# One-Step Selective Labeling of Native Cell-Surface Sialoglycans by Exogenous α2,8-Sialylation

**DOI:** 10.1101/2023.08.01.551455

**Authors:** Jonathan L. Babulic, Joshua M. Kofsky, Marie E. Boddington, Youjin Kim, Emmanuelle V. Leblanc, Sophie Emberley-Korkmaz, Che C. Colpitts, Chantelle J. Capicciotti

## Abstract

Exo-enzymatic glycan labeling strategies have emerged as versatile tools for efficient and selective installation of glycan terminal motifs onto live cell-surfaces. Through employing specific enzymes and nucleotide sugar probes, cells can be equipped with defined glyco-epitopes for modulating cell function or selective visualization and enrichment of glycoconjugates. Here, we identify *Campylobacter jejuni* sialyltransferase Cst-II I53S as a tool for cell-surface glycan modification, expanding the exo-enzymatic labeling toolkit to include installation of α2,8-disialyl epitopes. Labeling with Cst-II was achieved with biotin- and azide-tagged CMP-Neu5Ac derivatives on a model glycoprotein and on native sialylated cell-surface glycans across a panel of cell lines. The introduction of modified Neu5Ac derivatives onto cells by Cst-II was also retained on the surface for 6 h. By examining the specificity of Cst-II on cell surfaces, it was revealed that the α2,8-sialyltransferase primarily labeled N-glycans, with O-glycans labeled to a lesser extent, and there was an apparent preference for α2,3-linked sialosides. This approach thus broadens the scope of tools for selective exo-enzymatic labeling of native sialylated glycans and is highly amenable for construction of cell-based arrays.

## Introduction

The complex glycans and glycoconjugates displayed on cell surfaces play essential roles in many physiological and disease processes. Tools to interrogate the structures and functions of glycans and glycoconjugates in a native cellular environment are therefore of great interest to study their involvement in normal and pathological events.^1^ Metabolic oligosaccharide engineering (MOE) approaches have been used to incorporate monosaccharides bearing small bioorthogonal chemical reporters into cellular glycans, which can be selectively ligated with a complementary bioorthogonal probe for visualization or capture.^2, 3^ MOE relies on native cellular biosynthetic machinery, limiting the types of reporters that are tolerated throughout the labeling process. It also often results in broad, global incorporation of the monosaccharide probe into various glyco-epitopes and glycan subclasses, as selectivity is difficult to achieve.^4–6^ Complementary selective exo-enzymatic glycan labeling (SEEL) strategies have since emerged as valuable tools for efficient and specific labeling of cellular glycans.^7–9^ In SEEL, an exogenously administered glycosyltransferase is used to transfer and display nucleotide-sugar derivatives directly onto cell-surface glycans. By circumventing dependence on the native cellular biosynthetic pathways, SEEL is amenable to all cell types, can employ a broad range of probes, and results in efficient cell-surface glycan labeling with high glycosidic linkage and glycan subclass specificity.^7–10^ SEEL has thus garnered attention as a strategy to selectively install or edit specific glyco-epitopes and terminal motifs on cellular glycans.

One prominent application of SEEL is the selective sialylation of cellular glycans with sialyltransferases and CMP-sialic acid (CMP-Neu5Ac: cytidine-5’-monophospho-*N*-acetyl-neuraminic acid) donor substrates. Sialic acid (Neu5Ac) is typically found at the non-reducing terminal end of glycans, forming defined sialylated glyco-epitopes, as it can be attached through an α2,3- or α2,6-linkage to galactose (Gal) or *N-*acetylgalactosamine (GalNAc), or an α2,8-linkage to sialic acid.^11^ It is well established that the identity of the sialylated glyco-epitope plays distinct critical roles in diverse biological processes including cell adhesion, immune responses, cancer progression and pathogen interactions.^12^ Since sialic acids cap the termini of glycans, they are ideally suited for manipulation by SEEL, permitting the interrogation of the binding specificities of glycan-binding proteins and the study of the biological function of defined sialosides. For example, α2,3- and α2,6-sialyltransferases have been used for linkage-specific editing of cell-surface glycans with CMP-Neu5Ac to construct a live-cell array to study the contributions of these glyco-epitopes for in influenza A virus infection.^13^ The considerable substrate promiscuity of sialyltransferases has also been harnessed in SEEL using unnatural CMP-Neu5Ac derivatives bearing diverse chemical handles for visualization and enrichment of glycoconjugates, and cellular gain-of-function through introduction of functionally active biomolecules.^8, 10, 14–16^ The α2,6- sialyltransferase ST6Gal1 and α2,3-sialyltransferase ST3Gal1 have been employed to selectively label cellular N- or O-linked glycans, respectively, using CMP-Neu5Ac derivatives containing azides, alkynes, and other reporter probes.^7–9, 17–21^ Alkyne-linked Neu5Ac derivatives have also been selectively displayed on CHO cells using α2,3- and α2,6-sialyltransferases, where conjugation to a library of azide-bearing molecules enabled the discovery of epitope-specific high-affinity Siglec-15 ligands.^20^

While considerable advances have been made employing α2,3- and α2,6-sialyltransferases for cell-surface glycan editing through SEEL, α2,8-linked sialylation remains relatively unexplored.^7–10, 13, 17^ We were therefore interested in expanding the enzyme repertoire for selective exo-enzymatic editing to install α2,8-disialyl motifs onto cell-surface glycans. The α2,8-disialyl glyco-epitope is found as a common component of glycoproteins and gangliosides, such as the disialoganglioside GD2, which is overexpressed in neuroblastoma, osteosarcoma, and many other cancer types, and is a promising target for cancer immunotherapy.^22–26^ The α2,8-disialyl motif (along with the bis-sialylated Core 1 O-glycan) has also been implicated as an important ligand for Siglec-7, as well as other Siglecs that have been identified as glyco-immune checkpoint molecules and candidate targets for immune checkpoint blockade.^27–41^ Disialosides can also be further extended into α2,8-linked polysialic acid, which has diverse roles in neurobiology and cancer metastasis.^42–45^

In humans, α2,8-disialyl motifs are primarily biosynthesized by ST8Sia1 on glycolipids and ST8Sia6 on glycoproteins, but other ST8Sia enzymes are also implicated in some cases.^46–49^ The α2,8-disialyl motif can also be synthesized *in vitro* employing a mutant of the bifunctional sialyltransferase Cst-II from *Campylobacter jejuni*.^50^ This mutant Cst-II enzyme has a 32-amino acid truncation and I53S point mutation (Cst-IIΔ32^I53S^) that results in increased stability and enhanced α2,8-transferase activity. Cst-IIΔ32^I53S^ has been used in the synthesis of α2,8-disialyl epitopes on small oligosaccharides, complex glycan structures, and therapeutic glycoproteins, however its use for α2,8-sialylation on live cells has not been reported.^30, 50–58^

Here, we describe an approach for labeling native sialylated glycans through exogenous α2,8-disialylation with the *C. jejuni* sialyltransferase Cst-IIΔ32^I53S^ (hereafter referred to as Cst-II) (Fig. 1). We demonstrate that Cst-II can be used in a two-step labeling approach with an azide-conjugated CMP-Neu5Ac derivative, and in a direct one-step labeling using a biotinylated CMP-Neu5Ac, on both a model glycoprotein and live cell surfaces. We have found that Cst-II installs α2,8-linked sialic acid derivatives predominantly on N-linked glycans, but that this enzyme also labels O-glycans to a lesser extent, and it has an apparent general preference for installation onto α2,3-linked sialosides. Through this study, we expand the toolkit of cell-surface glycan engineering to exo-enzymatic installation of α2,8-disialyl motifs, while also providing a strategy for labeling native sialylated glycans and defining the substrate scope of the sialyltransferase Cst-II for cell-surface glycan editing.

**Figure 1.**
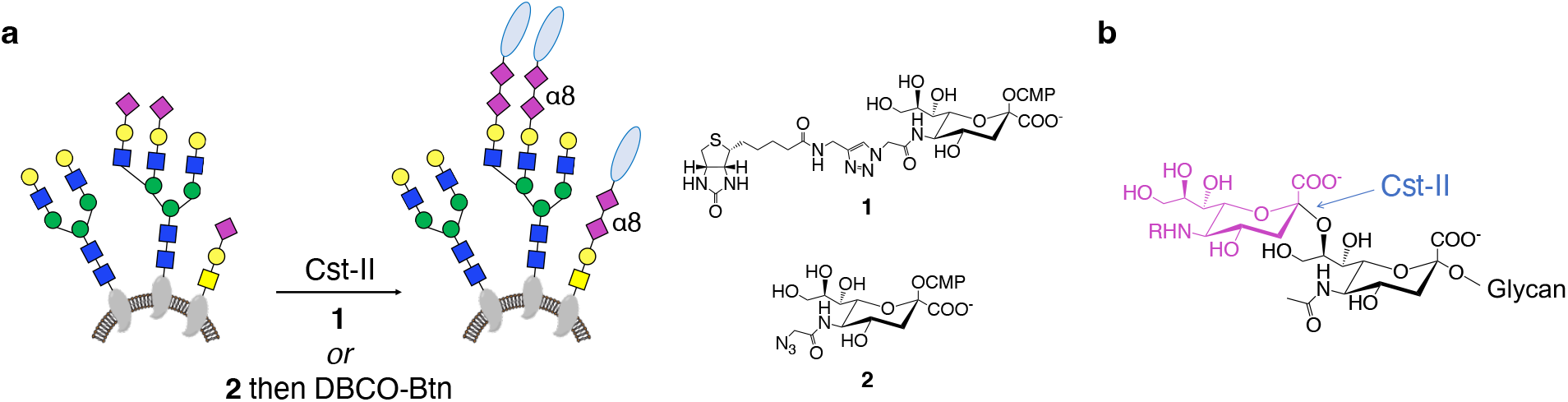
Cell-surface labeling through exo-enzymatic sialylation. **a)** One- and two-step labeling of native sialylated glycans through α2,8-sialylation with Cst-II and CMP-Neu5biotin (**1**) or CMP-Neu5Az (**2**). **b)** Structure of the α2,8-disialyl epitope produced by transfer of a CMP-Neu5Ac derivative with Cst-II.

## Results and Discussion

### Glycan labeling of fetuin

Cst-II has been used to install native CMP-Neu5Ac on fetuin (Fet) and other glycoproteins as an α2,8-disialyl epitope primer for polysialylation,^55, 59^ however it has been reported that a BODIPY-conjugated CMP-Neu5Ac is not tolerated for direct transfer onto Fet.^57^ A crystal structure of Cst-II with a CMP-3F-Neu5Ac analog suggests that the C-5 acetamido moiety of Neu5Ac is solvent-exposed.^60^ Thus, prior to labeling cellular glycans, we sought to examine the substrate tolerance of Cst-II with C5 azide-modified (**2**: CMP-Neu5Az) and biotin-modified (**1**: CMP-Neu5biotin) donors for enzymatic glycan labeling through α2,8-disialylation on the model glycoprotein Fet.

CMP-Neu5biotin (**1**) and CMP-Neu5Az (**2**) were prepared by previously reported protocols (Scheme S1, Supplementary Information).^8, 61^ The tolerance of Cst-II for the two CMP-Neu5Ac derivatives was then assessed by labeling fetuin by incubation with Cst-II and either CMP-Neu5biotin (**1**) in a one-step approach, or CMP-Neu5Az (**2**) which was detected by a strain-promoted azide-alkyne cycloaddition (SPAAC) reaction using DBCO-biotin (DBCO-Btn) in a two-step approach. Immunoblot with streptavidin demonstrated both substrates could successfully label Fet (Fig. 2a, Fig. S1) with the direct one-step labeling approach with CMP-Neu5biotin (**1**) showing stronger labeling, consistent with reports employing other sialyltransferases.^7, 8, 10^ To confirm Fet sialylation was a requirement for enzyme activity to form the labeled α2,8-disialylated product, Cst-II labeling with both **1** and **2** was repeated on asialofetuin (AFet). Labeling of AFet with **1** or **2** was barely detected above background by streptavidin Western blot analysis, demonstrating a dependence on sialylated substrates (Fig. 2).

**Figure 2.**
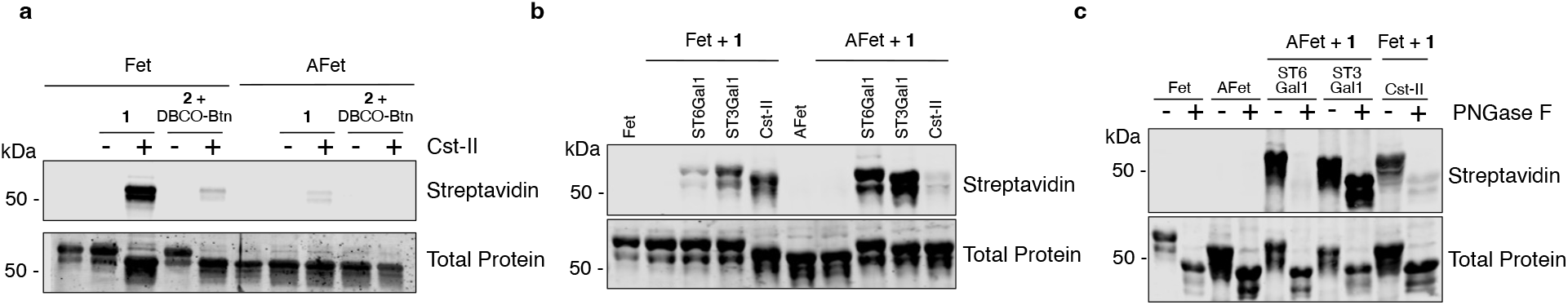
Enzymatic glycan labeling on fetuin. **a)** Bovine fetuin (Fet) or asialofetuin (AFet) was labeled with Cst-II and CMP-Neu5biotin (**1**) or CMP-Neu5Az (**2**) with subsequent conjugation to DBCO-s-biotin (DBCO-Btn) and immunoblotted with streptavidin-800CW. **b)** Fet or AFet were remodeled with **1** and ST6Gal1, ST3Gal1, or Cst-II. **c)** AFet labeled with **1** and ST6Gal1 or ST3Gal1, and Fet labeled with Cst-II, were treated with PNGase F prior to analysis by immunoblot with streptavidin-800CW.

Having demonstrated that Cst-II tolerates CMP-Neu5biotin (**1**) for a one-step transfer, we next examined if Cst-II had a preference for N- or O-linked glycans on the Fet substrate. Cst-II labeling was compared to the sialyltransferases ST6Gal1 and ST3Gal1, which selectively sialylate N- and O-linked glycans, respectively (Fig. 2).^21^ Both ST6Gal1 and ST3Gal1 successfully labeled Fet and AFet with **1**, with a greater intensity observed on the desialylated AFet substrate. Cst-II-labeled Fet and ST6Gal1- or ST3Gal1-labeled AFet were then treated with peptide N-glycosidase F (PNGase F) to release N-linked glycans. As expected, a complete loss of the biotin signal was observed when ST6Gal1-labeled AFet was treated with PNGase F. No change in signal was observed with ST3Gal1-labeled AFet, but N-glycan cleavage was apparent by a decrease in the observed molecular weight (Fig. 2).^17, 21^ The biotin signal was strongly decreased upon PNGase F treatment of Cst-II-labeled Fet, with a faint residual signal remaining. The extent of signal loss with PNGase F treatment suggests that labeling with Cst-II on Fet primarily occurred on N-glycans, with some transfer onto O-glycans. While the observed results may be due to a preference of Cst-II for N-linked glycan substrates, this could also be attributed to a greater abundance of terminal sialic acid residues reported to be on the N-glycans of Fet compared to O-glycans.^55, 62, 63^ Previous work using Cst-II on defined glycan substrates has also shown a broad substrate scope with oligosaccharides,^50, 51, 56, 60^ glycolipids,^53, 54^ α2,3- or α2,6-sialylated N-glycans,^30^ and N- and O-glycans on glycoproteins and glycopeptides.^55, 57, 59^

### Cell-surface glycan labeling

Next, we examined Cst-II for the labeling of native cell-surface sialoglycans to install the α2,8-disialyl epitope onto cells using CMP-Neu5Ac derivatives **1** and **2**. We selected a broad panel of human cell lines to investigate, including adherent embryonic kidney HEK293T/17 cells and breast cancer cell lines MCF-7, SK-BR-3, and HS-578-T, as well as suspension U937 monocytes and RAJI B cells. Initial labeling conditions were based on previous reports using other sialyltransferases;^8, 18, 50^ thus, cells were treated with 200 μg/mL Cst-II and 100 μM CMP-Neu5biotin (**1**) or CMP-Neu5Az (**2**) for 2 hours at 37°C. Display of **1** was detected by streptavidin staining followed by flow cytometry analysis, whereas the azide reporter of **2** was conjugated to DBCO-Btn via SPAAC prior to streptavidin flow cytometry detection.

In all cell lines examined, we observed that Cst-II tolerated the CMP-Neu5biotin (**1**) probe, demonstrating this enzyme is capable of labeling live cells with the α2,8-disialyl motif through a one-step SEEL approach (Fig. 3). The extent of labeling was variable across the various cell lines surveyed, suggesting different abundances of substrates for Cst-II across different cell lines. To examine whether increased labeling intensity was consistent with higher sialic acid-specific lectin binding, we probed each cell line with *Sambucus nigra* agglutinin (SNA) and *Maackia amurensis* lectin-II (MAL-II). SNA preferentially binds to α2,6-linked sialosides, whereas MAL-II recognizes α2,3-linked sialosides. Higher intensity labeling with Cst-II generally corresponded to higher sialic acid-binding lectin staining (Fig. S2). For example, HS-578-T cells had the highest degree of labeling with Cst-II and **1**, and these cells exhibited the highest binding to both SNA and MAL-II (Fig. 3, Fig. S2). However high lectin binding did not always correlate with a greater degree of Cst-II labelling as MCF-7 cells labeled weakly with Cst-II, and yet bound considerably to MAL-II. A two-step Cst-II labeling approach with CMP-Neu5Az (**2**) was also effective at labeling, but generally showed weaker labeling when compared to the one-step approach (Fig. 3), consistent with reports using other sialyltransferases and our studies on Fet (Fig. 2).^7, 8, 10, 16^

**Figure 3.**
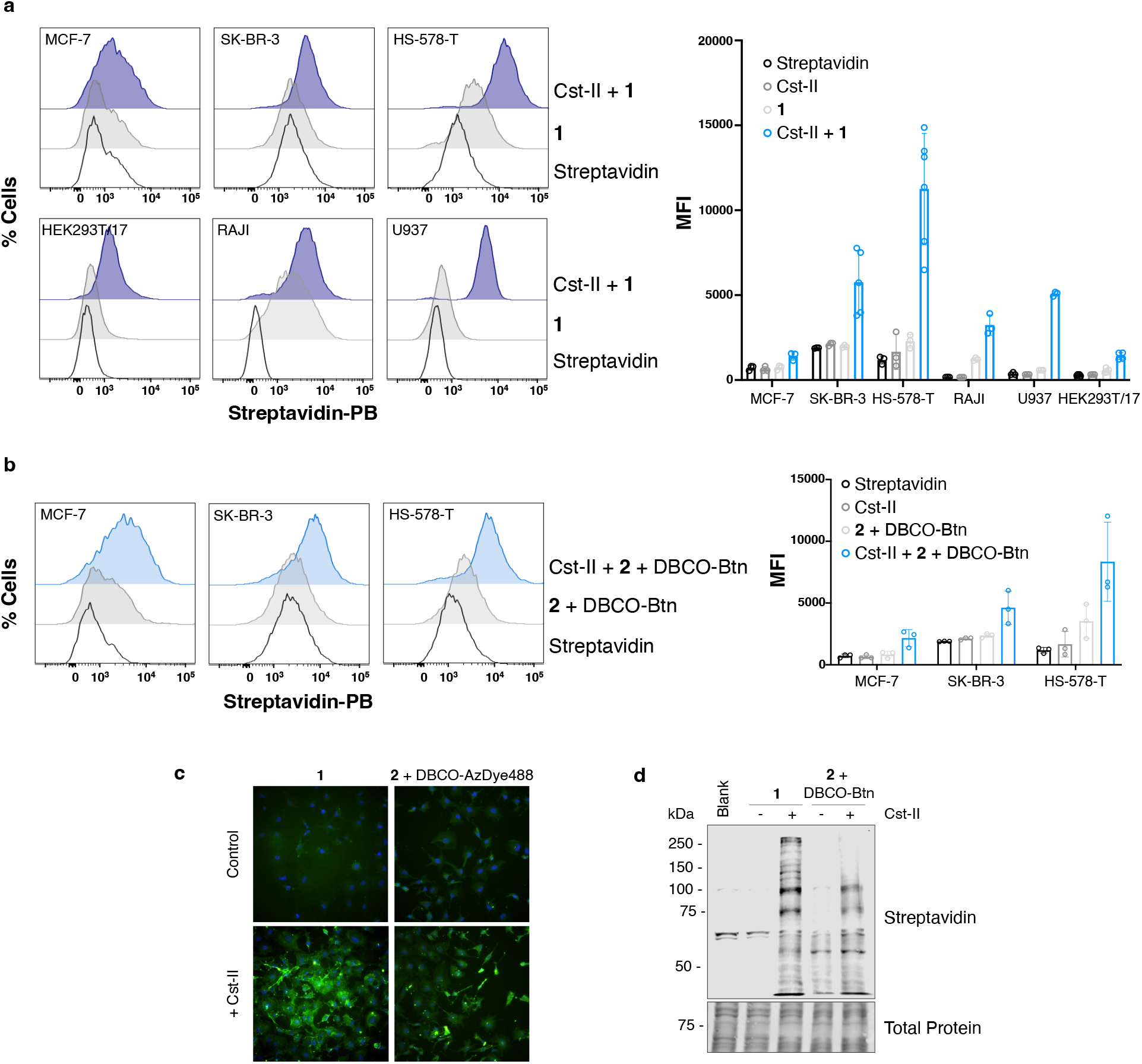
Cell-Surface Labeling with Cst-II. **a)** Cells were labeled with 200 μg/mL Cst-II and 100 μM CMP-Neu5biotin (**1**) for 2 h at 37 °C. Labeling with **1** on live cell-surfaces was assessed by flow cytometry. Cells were stained with Streptavidin-Pacific Blue (−PB) and co-stained with PI to exclude nonviable cells. Median fluorescence intensity (MFI) of each sample was calculated. Error bars represent the standard deviation of at least three replicates (n = 3). **b)** Cells were labeled with 200 μg/mL Cst-II and 100 μM CMP-Neu5Az (**2**) and subsequently tagged with 50 μM DBCO-Btn in a two-step protocol. Cells were stained with Streptavidin-PB and co-stained with PI to exclude nonviable cells to assess labeling by flow cytometry. Median fluorescence intensity (MFI) of each sample was calculated. Error bars represent the standard deviation of at least three replicates (n = 3). **c)** Fluorescence microscopy of HS-578-T cells labeled with Cst-II. **d)** Detection of glycoconjugates labeled on HS-578-T cells by one- or two-step cell-surface remodeling by immunoblot with streptavidin-800CW.

HS-578-T cells, which showed the most efficient labeling, were next used to further optimize the SEEL strategy with Cst-II. We confirmed that incubation with Cst-II alone did not alter binding of SNA or MAL-II, indicating that the Cst-II mutant used in our studies was not desialylating cells, activity that has previously been reported on substrates *in vitro* (Fig. S3).^50^ Cells labeled with CMP-Neu5biotin (**1**) or CMP-Neu5Az (**2**) were also imaged by fluorescence microscopy, confirming the introduction of both probes onto cell surfaces (Fig. 3c). Furthermore, examination of cell lysates by immunoblot showed that a broad range of glycoproteins were labeled by Cst-II (Fig. 3d). Optimization of the SEEL conditions using Cst-II demonstrated that the ideal concentration of Cst-II was 200 μg/mL and reaction time was 2 h to achieve the highest level of labeling (Fig. S4a,b). Titration of **1** showed that transfer of the probe was optimal at 100 μM and did not significantly increase with higher donor concentrations (Fig. S4c).

We also examined the lifetime of the biotin probe display after SEEL with Cst-II and **1** by culturing labeled HS-578-T cells in complete medium and detecting biotinylation by flow cytometry and Western blotting. Analysis by flow cytometry showed that streptavidin binding was only slightly decreased 6 h post-labeling with **1** and Cst-II (Fig. 4a), whereas Western blotting showed a disappearance of many biotin-tagged glycoproteins compared to immediately post-labeling (*t* = 0 h; Fig. 4b). A marked decrease in signal was observed 24 h post-labeling with **1** and Cst-II by both flow cytometry and Western blot analysis (Fig. 4). Weak labeling was detected 72 h post-labeling with **1** and Cst-II by flow cytometry, with a few persistent biotinylated glycoprotein bands present by Western blot. These results suggest that a small sub-population of sialylated glycoconjugates that were labeled with the α2,8-disialyl glyco-epitope by Cst-II are turned over more slowly, resulting in a longer display of the Neu5Ac derivative. This persistent labeling after SEEL has previously been observed with the sialyltransferase ST6Gal1.^10^ Notably, the retained signal was stronger with ST6Gal1 24 and 72 h post-labeling, which is likely attributed to ST6Gal1 being a more efficient enzyme for SEEL compared to Cst-II. Persistent cell-surface display of synthetic ligands is important for applying cell-surface engineering towards biological studies or development of cell-based therapies, and our findings suggest that Cst-II introduction of a modified Neu5Ac derivative on cell-surfaces can be retained for at least 6 h.^10, 64, 65^

**Figure 4.**
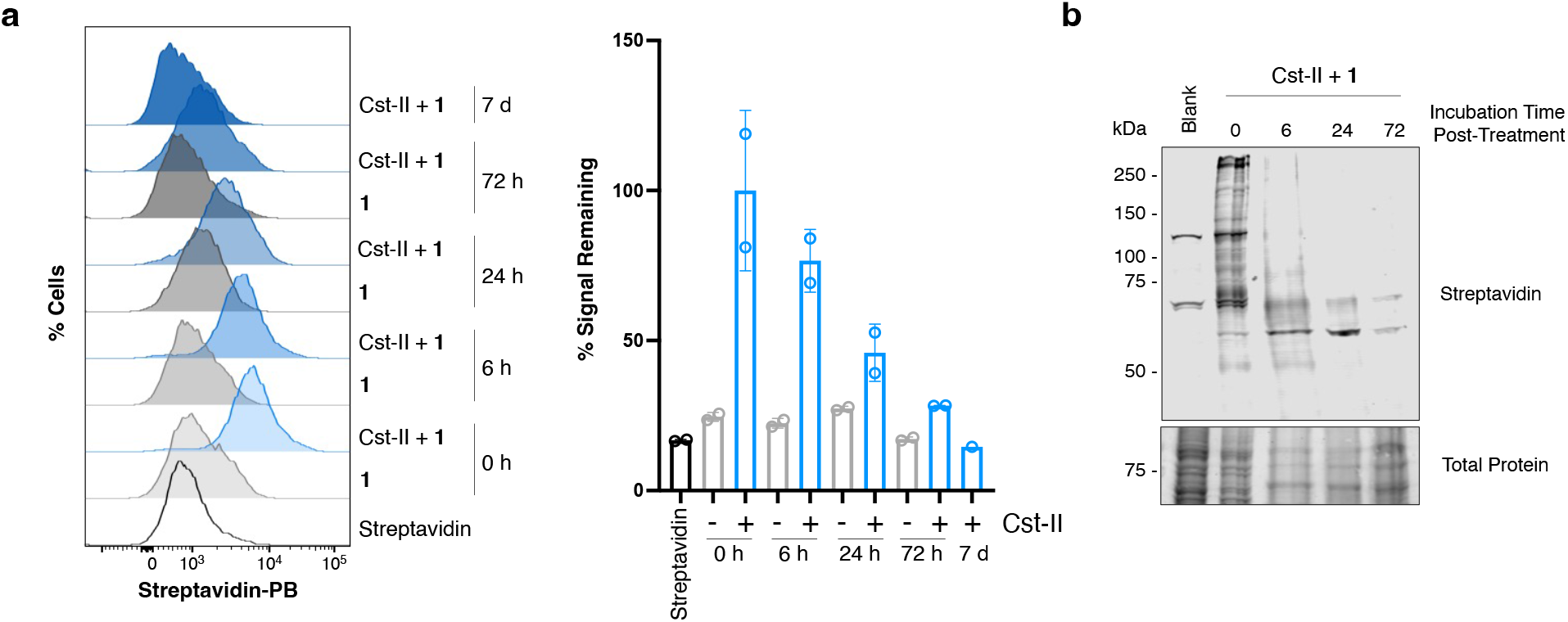
Timeline of display of 1 on cell surfaces with Cst-II. HS-578-T cells were incubated with 100 μM of CMP-Neu5biotin (**1**) and with or without 200 μg/mL Cst-II for 2 hours at 37 °C. Cells were then washed and incubated in complete medium for the timepoints indicated. **a)** Cells were stained with Streptavidin-Pacific Blue, co-stained with PI to exclude nonviable cells, and labeling was assessed by flow cytometry. Median fluorescence intensity (MFI) of each sample was calculated. Error bars represent the standard deviation of two replicates (n = 2). **b)** Cells were lysed and labeling with **1** was assessed by immunoblot with streptavidin-800CW.

### Cst-II labels N- and O-linked glycans

Having confirmed that Cst-II is able to label cell-surface glycans, we were interested in exploring which subclasses of glycans were substrates for Cst-II. Our results from SEEL of Fet with **1** and Cst-II suggested predominant labeling of N-glycans and weak labeling of O-glycans (Fig. 2). To examine this specificity in the context of cell-surface SEEL, we treated HS-578-T cells for 72 h with common inhibitors of major glycan biosynthesis pathways prior to labeling with Cst-II and CMP-Neu5biotin (**1**). Treatment with kifunensine, an inhibitor of complex N-glycan biosynthesis, greatly reduced labeling with Cst-II on HS-578-T cells, suggesting the majority of labeling is occurring on complex N-glycans (Fig. 5a). Benzyl-α-GalNAc treatment to inhibit O-glycan biosynthesis also significantly reduced Cst-II labeling, but to a lesser extent than kifunensine, suggesting that O-glycans are also exo-enzymatically labeled by Cst-II. However, blockade of glycolipid biosynthesis with GENZ-123346 did not significantly alter labeling with Cst-II, indicating that glycolipids are not significant substrates for Cst-II at the cell surface under these labeling conditions. This may be due to the specificity of Cst-II or may be related to the poor accessibility of glycolipids at the cell-surface as Cst-II has been successfully used to synthesize α2,8-disialylated glycolipid glycans *in vitro*.^53, 54, 58^

**Figure 5.**
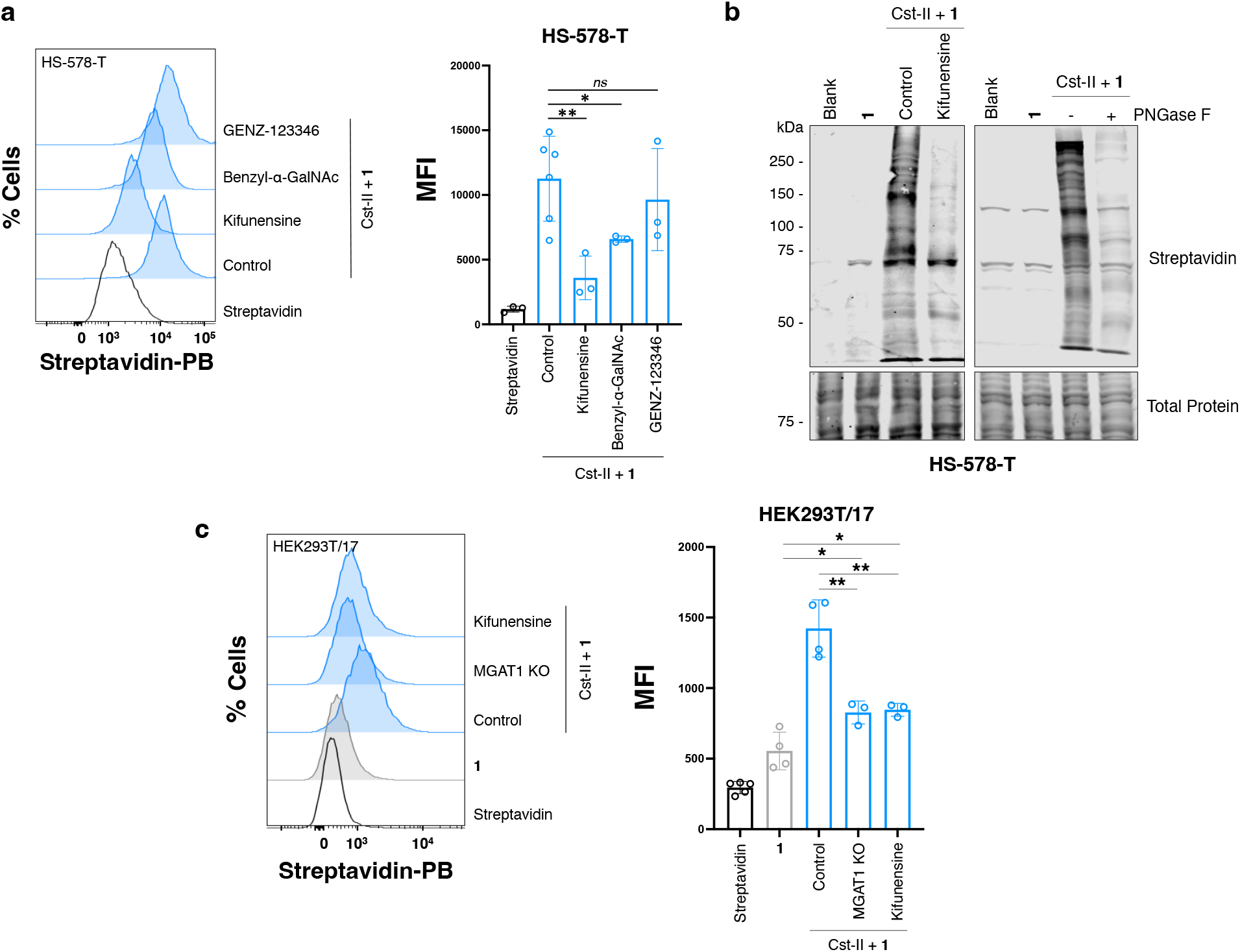
Exploring the glycan substrates of Cst-II. **a)** HS-578-T cells were cultured for 72 h with inhibitors of N-glycan (kifunensine), O-glycan (benzyl-α-GalNAc), or glycolipid (GENZ-123346) biosynthesis. Cells were then labeled with Cst-II and 100 μM CMP-Neu5biotin (**1**). Labeling with **1** on live-cell surfaces was assessed by flow cytometry. Cells were stained with Streptavidin-Pacific Blue and co-stained with PI to exclude nonviable cells. Median fluorescence intensity (MFI) of each sample was calculated. Error bars represent the standard deviation of at least three replicates (n = 3). Statistical significance was assessed by unpaired t-test with *ns* = not significant, *p < 0.05, and **p < 0.01. **b)** Cells were labeled with Cst-II and CMP-Neu5biotin (**1**) with or without treatment with kifunensine, or lysed, incubated with or without PNGase F. Lysates were analyzed by immunoblot and labeling with **1** was detected with streptavidin-800CW. **c)** HEK293T/17 cells cultured with or without kifunensine for 72 h and HEK293T/17 MGAT1 KO cells were labeled with Cst-II and 100 μM CMP-Neu5biotin (**1**). Labeling with **1** on live-cell surfaces was assessed by flow cytometry. Cells were stained with Streptavidin-Pacific Blue and were co-stained with PI to exclude nonviable cells.

To confirm the cell-surface substrate specificity of Cst-II, lysates of HS-578-T cells labeled with **1** and Cst-II were incubated with PNGase F and probed by streptavidin immunoblot (Fig. 5b). As observed with fetuin, labeling was greatly decreased by PNGase F treatment, but was not completely abolished, suggesting predominant labeling of N-glycans and weaker labeling of O-linked glycans. Furthermore, knock out (KO) of MGAT1 (Fig. S5), an enzyme essential for the maturation of complex-type N-glycans, by CRISPR/Cas9 in HEK293T/17 cells validated the preference of Cst-II for cell-surface N-glycans (Fig. 5c). Cst-II SEEL of HEK293T/17-MGAT1 KO cells with **1** dramatically decreased labeling to a similar level as HEK293T/17-WT cells treated with kifunensine (Fig. 5c). Consistent with what was observed with HS-578-T cells, labeling levels on HEK293T/17-MGAT1 KO cells did not decrease to the enzyme-free control, suggesting that sialylated O-glycans are being labeled on HEK293T/17 cells.

### Cst-II labels α2,3- and α2,6-linked sialosides

In humans, sialylation can occur through α2,3- and α2,6-linkages to an underlying Gal or GalNAc, or α2,8-linkages to other sialosides. While Cst-II has been shown to tolerate both α2,3- and α2,6-sialosides on Gal *in vitro*,^30, 50, 53, 56, 58^ given the range in labeling efficiency (Fig. 3) and the variable lectin binding data associated with different cell lines (Fig. S2), we were interested in determining if we could probe the specific sialylated epitopes on cell surfaces that were serving as substrates for Cst-II. To assess whether Cst-II displays specificity for the underlying sialoside glycosidic linkage, prior to labeling cells with **1** and Cst-II, cells were pre-treated with either an α2,3-specific sialidase from *Streptococcus pneumoniae* (Neu-S), or a broadly acting (α2,3/6/8) sialidase from *C. perfringens* (NanH). The α2,3-selectivity of Neu-S was assessed and confirmed by lectin staining with SNA and MAL-II, showing that as expected, only MAL-II binding decreased upon Neu-S treatment (Fig. S6).

The Neu-S or NanH pre-treatments revealed differences in Cst-II labeling in three different cell lines. After pre-treatment with the broadly acting sialidase NanH, labeling with Cst-II was abolished (Fig. 6a). In contrast, labeling was significantly reduced, but not eliminated, after pre-treatment with α2,3-specific sialidase Neu-S. These results indicate that both α2,3- and α2,6-linked sialosides are substrates for Cst-II on cell surfaces. In these sialidase pre-treatment studies, we noticed an interesting observation with the Cst-II-free controls (treatment with **1** alone) on SK-BR-3 and HS-578-T cells. After pre-treatment of these cells with either NanH or Neu-S, labeling was increased in the presence of **1** alone (no Cst-II enzyme) compared to the natively sialylated cells treated with **1** alone. This is likely due to transfer of **1** by endogenous sialyltransferases to the exposed LacNAc acceptor sites after sialidase treatment, which has previously been reported with CMP-Neu5Ac derivatives on B-cell lines and other immune cells.^18, 66–69^ Notably, there was no statistical difference in the extent of endogenous sialyltransferase labeling with **1** after either NanH or Neu-S pre-treatment in SK-BR-3 or HS-578-T cells. MCF-7 cells do not appear to have this endogenous sialyltransferase activity as labeling with **1** alone was not increased above background levels (Fig. 6).

**Figure 6.**
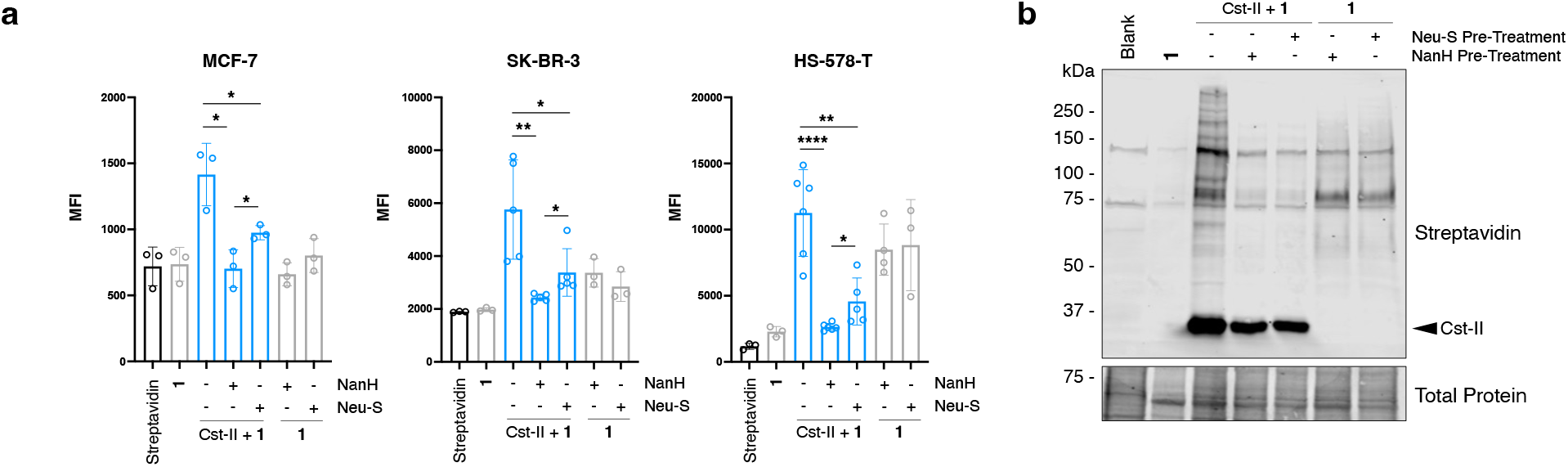
Cst-II labeling on sialidase pre-treated cells. **a)** Cells were treated first pre-treated with or without broadly acting sialidase NanH or α2,3-specific sialidase Neu-S and then labeled with Cst-II and **1** or **1** alone. Labeling with **1** on live-cell surfaces was assessed by flow cytometry. Cells were stained with Streptavidin-Pacific Blue and were co-stained with PI to exclude nonviable cells. Median fluorescence intensity (MFI) of each sample was calculated. Error bars represent the standard deviation of at least three replicates (n = 3). Statistical significance was assessed by unpaired t-test with *ns* = not significant, *p < 0.05, **p < 0.01, and ****p < 0.0001. **b)** HS-578-T cells were pre-treated with or without NanH or Neu-S and then labeled and lysed to be resolved by immunoblot with streptavidin-800CW.

A streptavidin immunoblot of HS-578-T cell lysates showed that a distinct subset of glycoproteins are labeled with **1** by endogenous sialyltransferases post-sialidase treatment compared to labeling natively sialylated cells with Cst-II (Fig. 6b). The observation that endogenous extracellular sialyltransferase activity is present in some breast cancer cell lines is intriguing, notably because of emerging cancer immunotherapies employing methods to selectively desialylate tumours to activate immune cell killing.^33, 70, 71^ Recent work by Lau et al. found that extracellular ST6Gal1 present in cancer exosomes promoted breast cancer cell growth and invasiveness.^72^ While the source of CMP-Neu5Ac substrate for cancer cell extracellular ST6Gal1 has not been explicitly identified, platelets have been shown to supply CMP-Neu5Ac for exogenous sialyltransferases after inflammatory challenges.^72–74^ Further work is warranted to further investigate and characterize this extracellular sialyltransferase activity and its functional effects in cancer and other pathological states.

By flow cytometry streptavidin staining, it is evident that pre-treatment with NanH abolished labeling with **1** and Cst-II and that a low level of labeling is retained after α2,3-specific Neu-S pre-treatment. We were surprised to observe that for different sialidase pre-treatments on certain cell lines, labeling with **1** by endogenous sialyltransferases (the Cst-II-free control) was higher than when Cst-II was present (*e.g.* HS-578-T cells; Fig. 6a). We hypothesized that when sialylated acceptor sites were cleaved, the addition of Cst-II may be competing with endogenous sialyltransferase labeling activity by hydrolyzing **1**. Cst-II has been reported to have weak hydrolytic activity towards CMP-Neu5Ac,^60^ thus we investigated whether the enzyme hydrolyzed the CMP-Neu5biotin (**1**) probe. However, *in vitro* incubation of **1** with Cst-II for 2 h, the incubation time used for cell labeling, showed minimal hydrolysis by NMR analysis (Fig. S7), suggesting an alternate method of sequestering CMP-Neu5biotin from endogenous sialyltransferases.

Streptavidin immunoblot of HS-578-T lysates from cells treated in the presence of Cst-II and **1** showed a prominent biotin-labeled band at ~30 kDa, the approximate molecular weight of Cst-II (Fig. 6b). This band at 30 kDa was also visible in fetuin samples remodeled with Cst-II and **1** or **2**, but not when other sialyltransferases were used to label Fet (Fig. S1). We therefore postulated that Cst-II may be biotinylated with **1**, and after an overnight incubation of Cst-II alone with CMP-Neu5biotin (**1**), immunoblot showed a streptavidin-positive band for Cst-II (Fig. S8). This finding was surprising, however it provides an explanation for the decreased labeling of desialylated cells with **1** in the presence of Cst-II, and we hypothesize self-labeling of Cst-II may be outcompeting the endogenous sialyltransferases on cell-surfaces for the donor substrate **1** (Fig. 6).

While recombinant proteins expressed in *E. coli* are not typically glycosylated with sialylated glycans without extensive genetic engineering,^75, 76^ we sought to confirm that the labeling of Cst-II with **1** was not occurring due to an unanticipated glycosylation site. Pre-treatment of Cst-II with either sialidase (NanH) or galactosidase (BgaA) did not alter labeling of Cst-II (Fig. S8a), suggesting that it does not self-label through a sialic acid or galactose substrate. Additionally, no reduction in MW of the biotinylated-Cst-II band was observed in the PNGase F treated Cst-II-labeled Fetuin samples (Fig. S1). We also treated the enzyme with a mild sodium periodate oxidation to oxidize/cleave and remove the C-8 hydroxyl group of any potential sialic acids.^77^ While incubation of **1** with periodate treated Cst-II showed decreased labeling compared to untreated Cst-II (Fig. S8b), a reduction of Fet labeling using periodate-treated Cst-II was also observed. Since glycosidase treatments did not alter Cst-II self-labeling, the decrease observed after periodate treatment is likely due to a loss of enzyme activity, rather than due to loss of a glycosyl acceptor substrate for transfer of **1** onto Cst-II.

Generally, the results comparing Neu-S or NanH pre-treated cells indicate that Cst-II labels α2,3- and α2,6-linked sialosides, aligning with reports that both have served as substrates for Cst-II in chemoenzymatic glycan synthesis (Fig. 6a).^30^ Initial observations suggest that α2,3-sialosides may be the predominant substrate on cell surfaces (Fig. 6a). However, given that endogenous sialyltransferases on certain cell lines (SK-BR-3, HS-578-T; Fig. 6a/b) may be competing with exogenously introduced Cst-II for labeling desialylated cells, further detailed validation is needed to confirm this preference. Additionally, while Cst-II can label native sialosides on glycoproteins and cell-surfaces with an α2,8-disialyl epitope, it is evident that the Cst-II enzyme self-labels in the presence of **1** or **2** (Fig. 6; Fig. S1, S8). This observation may explain why cell-surface labeling with Cst-II is not as robust as other sialyltransferases such as ST6Gal1 – the enzyme may be sequestering the donor substrate by the formation of an unusual covalent product between the modified CMP-Neu5Ac donor (**1**) and Cst-II. We hypothesize this covalent enzyme-substrate adduct may be occurring through Tyr156 or Tyr162 in the active site. These residues are thought to aid in the generation of an oxocarbenium-like transition state of the donor substrate based on crystallization studies of Cst-II with CMP-3FNeu5Ac.^51, 60^ Furthermore, these residues, particularly Tyr156, may be close enough in proximity to act as a nucleophile (Fig. S9). Covalent enzyme-substrate complexes can occur in sialidase mechanisms, and while our data suggests the Cst-II mutant used in our studies is not acting as a sialidase or hydrolyzing the donor (Fig. S6, S7), formation of a covalent intermediate would provide an explanation for our observations.^78^ Full characterization of the Cst-II-**1** adduct is ongoing and may provide useful insight into possible mutations that could be made in the active site to disfavour self-reactivity and enhance cell-surface labeling using Cst-II.

## Conclusions

Selective exo-enzymatic cell-surface glycan labeling has previously been applied through α2,3- and α2,6-sialylation to label and functionalize galactose-capped glycoconjugates.^7–10, 13, 16, 17, 57^ Here, we report selective exo-enzymatic cell-surface labeling of sialoglycans through α2,8-sialylation by harnessing the sialyltransferase Cst-II from *C. jejuni*. The strategy is broadly applicable to different cell types and our results suggest that Cst-II labels sialylated N- and O-linked glycans, but that there is a preference for N-glycan substrates. A one-step SEEL with Cst-II permits direct labeling of native sialylated glycoconjugates through α2,8-disialylation and represents a novel exo-enzymatic tool for probing the cell-surface sialome and displaying unnatural CMP-Neu5Ac derivatives. Furthermore, the use of a bacterial sialyltransferase makes the method accessible as this enzyme is simple to express and purify with high yields. We anticipate this strategy will be relevant for use with nucleotide-sugar derivatives bearing other functional groups, including photo-cross-linking probes or complex synthetic ligands, particularly as we have demonstrated that labeling by Cst-II can persist on cell surfaces for at least 6 h. The use of photo-cross-linking probes would facilitate the identification of glycan-binding proteins that may use the α2,8-disialyl epitope for recognition and biological functions, but have yet to be identified.^18, 66^ Appending complex synthetic ligands to the CMP-Neu5Ac donor derivative could expand applications of this SEEL strategy to cell-based glycan arrays, enabling the identification of high-affinity ligands for glycan-binding proteins or the development of cell-based therapies.^10, 13, 15, 18, 66, 79^ Furthermore, this approach may be useful in tandem with other glycosyltransferases for multiplex imaging of various cell-surface glycan epitopes with different nucleotide-sugar probes, such as through chemo-enzymatic histology.^19^ This work expands the toolkit of glycosyltransferases for applications in cell-surface glycan engineering will open up new avenues for the study of glycoconjugates and the biologically important α2,8-sialylated glycan epitopes.

## Associated Content

Supporting Information: Supplementary figures of flow cytometry and immunoblot studies; materials and methods for biological procedures and synthesis and characterization data.

## Acknowledgements

The authors thank Dr. Matthew Macauley (University of Alberta) for providing HS-578-T and U937 cell lines, and the plasmid encoding the gene for Neu-S. The authors also thank Dr. David Kwan (Concordia University) for providing the plasmid encoding the gene for BgaA. We acknowledge the New Frontiers in Research Fund (SSHRC-NFRF), the National Sciences and Engineering Research Council of Canada (NSERC) and the Canada Foundation for Innovation (CFI) for funding supporting this work. J.L.B. and M.E.B acknowledge NSERC for a CGS-D scholarship. J.M.K. and E.V.L. are supported by Vanier Canada Graduate Scholarships and Y.K. acknowledge NSERC for a PGS-D scholarship.

## Notes

### Competing Interest Statement

The authors have declared no competing interest.

