## Supporting Information for "One-Step Selective Labeling of Native Cell-Surface Sialoglycans by Exogenous α2,8-Sialylation"

### Contents

Supplemental Figures ........................................................................... S2
Biological Methods ............................................................................... S8
Chemo-Enzymatic Synthesis ................................................................. S15
NMR Spectra ........................................................................................ S18
References ............................................................................................ S21

### 1. Supplemental Figures

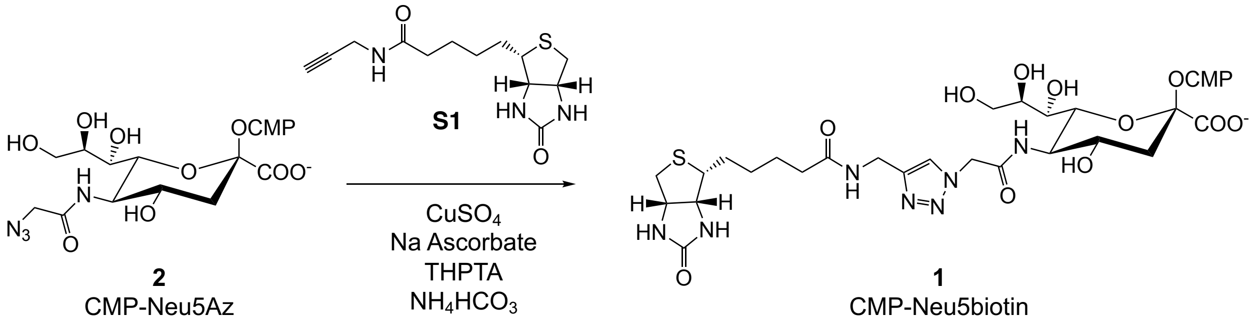


**Scheme S1.** Synthesis of CMP-Neu5biotin (**1**) by copper-catalyzed azide-alkyne cycloaddition (CuAAC) using donor and CMP-Neu5Az (**2**).

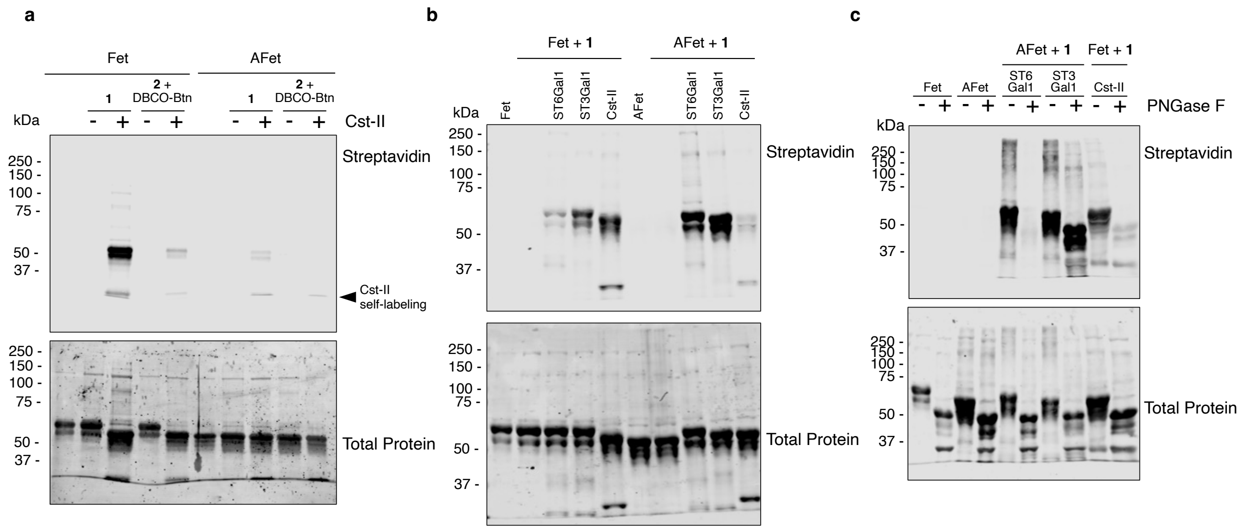


**Figure S1. Full length immunoblots of fetuin glycan labeling. a)** Bovine fetuin (Fet) or asialofetuin (AFet) was labeled with Cst-II and CMP-Neu5biotin (**1**) or CMP-Neu5Az (**2**) and subsequent conjugation to DBCO-biotin (DBCO-Btn) and immunoblotted with streptavidin-800CW. **b)** Fet or AFet was remodeled with **1** and ST6Gal1, ST3Gal1, or Cst-II. **c)** AFet labeled with **1** and ST6Gal1 or ST3Gal1 and fetuin labeled with Cst-II were treated with PNGase F prior to analysis by immunoblot with streptavidin-800CW.

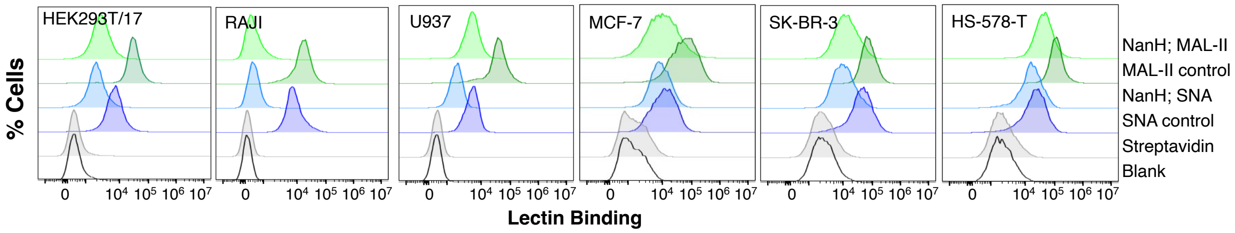


**Figure S2. Probing native sialylation patterns by lectin staining.** Cells were incubated with or without 50 μg/mL *C. perfringens* sialidase NanH for 30 min and subsequently stained with biotinylated SNA or MAL-II. Cells were then stained with Streptavidin-Pacific Blue and co-stained with PI to exclude nonviable cells and lectin binding was assessed by flow cytometry.

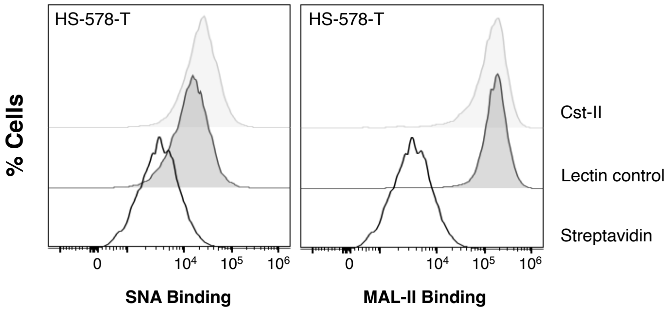


**Figure S3. Lectin binding to cells treated with Cst-II alone.** HS-578-T cells were treated with 200 μg/mL Cst-II for 2 hours at 37 °C and then stained with biotinylated SNA or MAL-II. Cells were then stained with Streptavidin-Pacific Blue and co-stained with PI to exclude nonviable cells and lectin binding was assessed by flow cytometry.

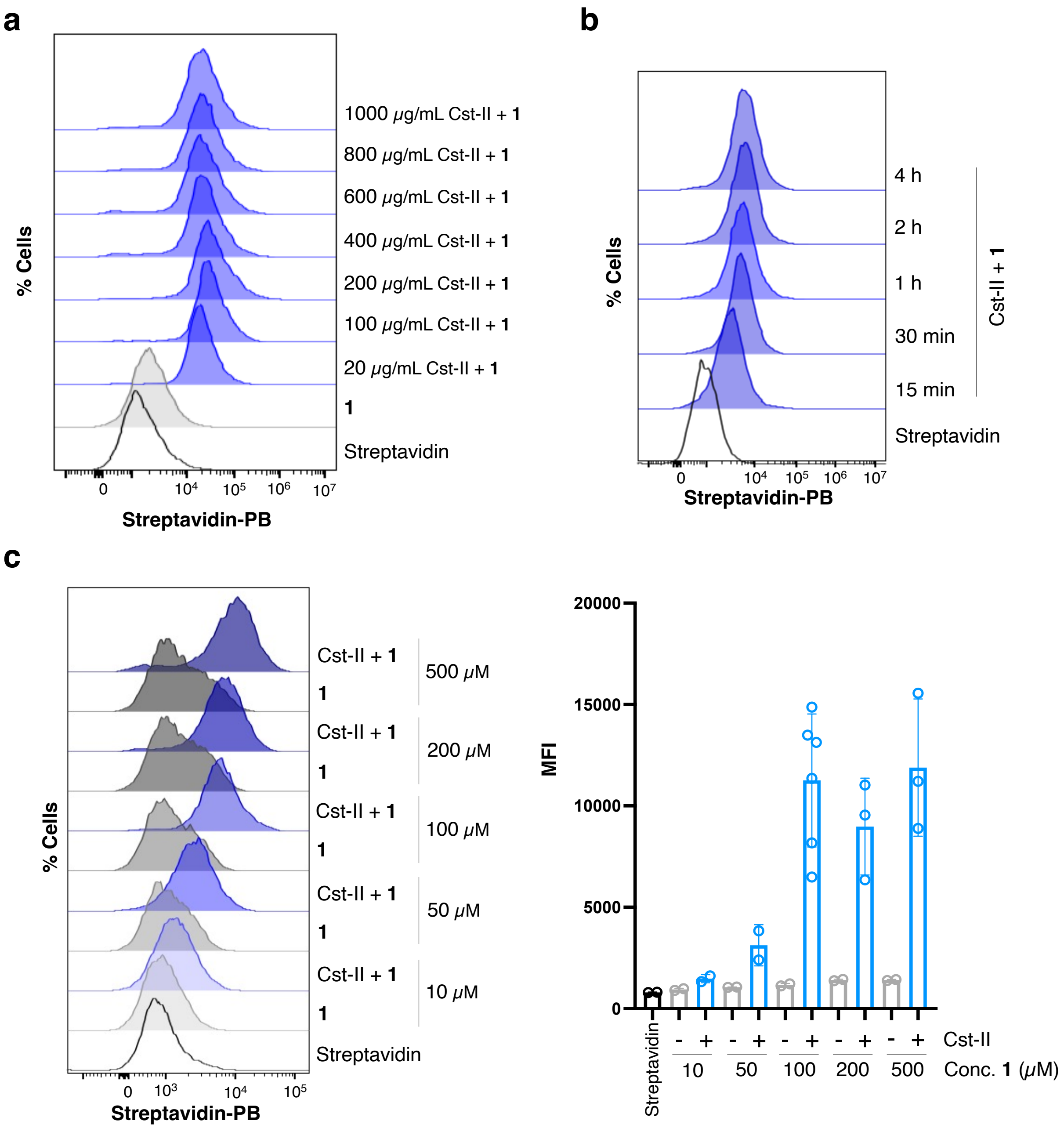


**Figure S4. Cell-surface display of CMP-Neu5biotin with Cst-II. a)** HS-578-T cells were incubated with 100 µM of CMP-Neu5biotin (**1**) and varying concentrations of Cst-II for 2 hours at 37 °C. Cells were then stained with Streptavidin-Pacific Blue and co-stained with PI to exclude nonviable cells and labeling was assessed by flow cytometry. **b)** Cells were incubated with 100 µM of CMP-Neu5biotin (**1**) and 200 µg/mL Cst-II. Labeling was assessed at various incubation timepoints by staining with Streptavidin-Pacific Blue. Cells were co-stained with PI to exclude nonviable cells and labeling was assessed by flow cytometry. **c)** Cells were incubated with varying concentrations of CMP-Neu5biotin (**1**) and 200 µg/mL Cst-II for 2 hours at 37 °C. Cells were then stained with Streptavidin-Pacific Blue and co-stained with PI to exclude nonviable cells and labeling was assessed by flow cytometry. Error bars represent the standard deviation.

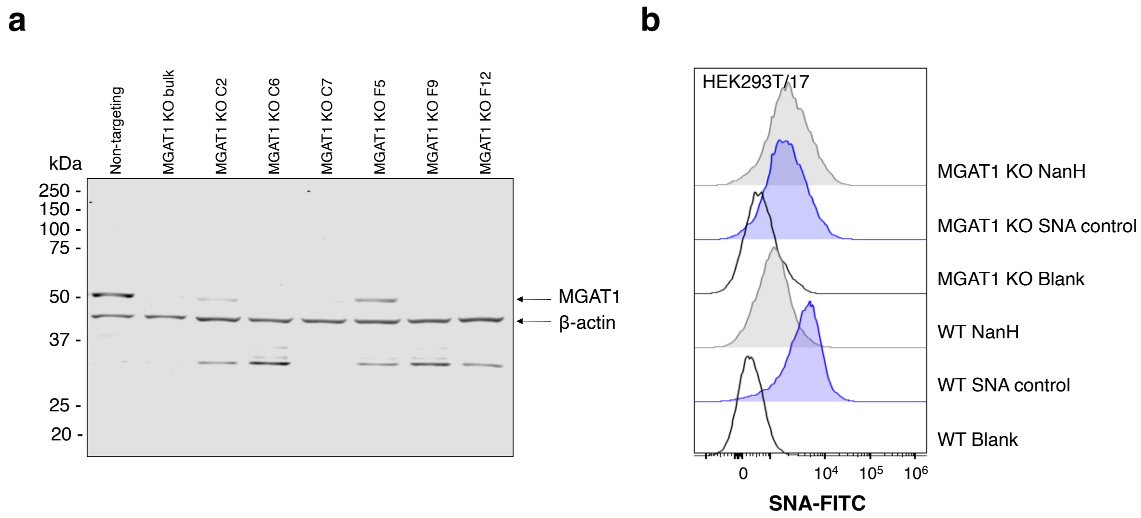


**Figure S5. Screening of MGAT1 KO cells. a)** HEK293T/17-MGAT1 KO clonal populations were immunoblotted with an anti-MGAT1 antibody and anti-β-actin as a loading control. Clone F9 was selected for subsequent experiments. **b)** Wild type HEK293T/17 cells or MGAT1 KO cells were incubated with or without 50 µg/mL *C. perfringens* sialidase NanH for 30 min and subsequently stained with 20 µg/mL SNA-FITC. Cells were co-stained with PI to exclude nonviable cells and assessed by flow cytometry.

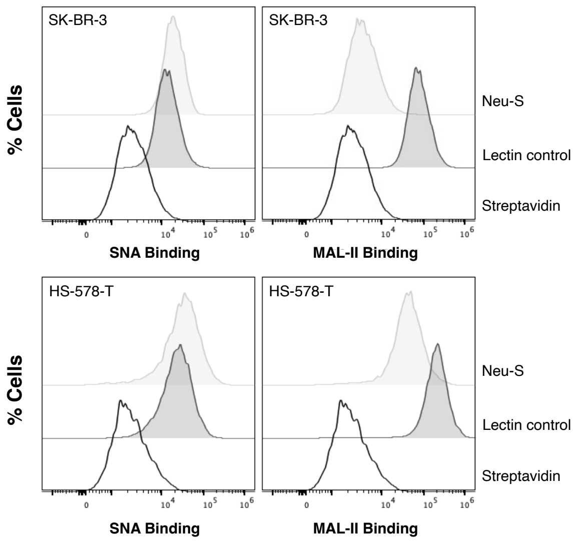


**Figure S6. Neu-S selectively releases α2,3-sialosides on cell-surfaces.** SK-BR-3 and HS-578-T cells were incubated with or without 50 µg/mL sialidase Neu-S for 30 min and subsequently stained with 20 µg/mL biotinylated SNA or MAL-II. Cells were then stained with streptavidin-Pacific Blue, co-stained with PI to exclude nonviable cells, and lectin binding was assessed by flow cytometry.

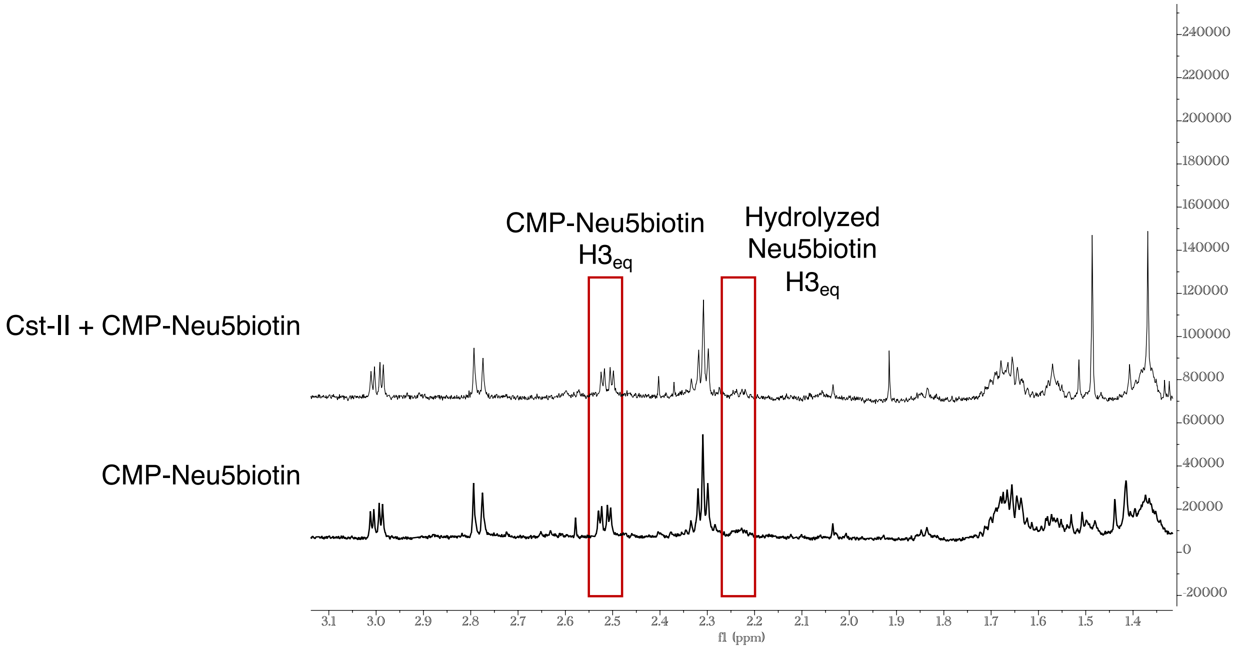


**Figure S7. Monitoring Cst-II incubation with CMP-Neu5Biotin by NMR.** CMP-Neu5biotin (**1**) was dissolved in Tris-HCl buffer (100 mM, pH 9) to a final concentration of 10 mM, with 20 mM $MgCl_2$. Cst-II (100 µg/ml) was added to the reaction tube and the reaction was incubated at 37°C, shaking at 180 rpm. After shaking for 2 h the enzyme was removed by centrifugation using an Amicon Ultra-10 (MWCO-10k) centrifugal filter. The filtrate was lyophilized and analyzed by NMR (700 MHz, $D_2O$ referenced to 4.79 ppm). CMP-Neu5biotin incubated with Cst-II for 2 h (top) was compared to a standard of untreated CMP-Neu5biotin (bottom). The $H3_{eq}$ peaks are highlighted, indicating minimal hydrolysis of the CMP-Neu5Ac sugar.

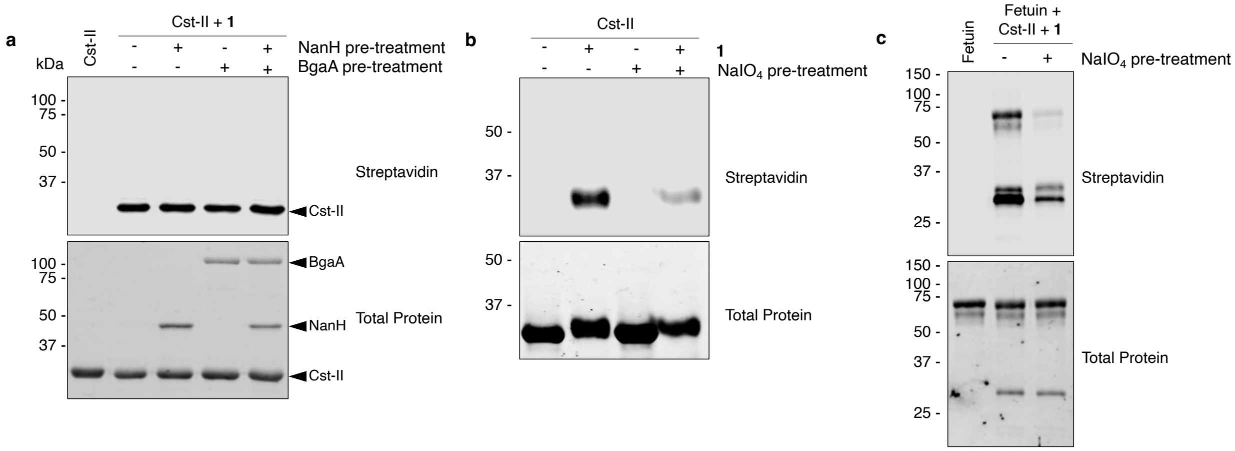


**Figure S8. Cst-II self-labeling with CMP-Neu5biotin. a)** Cst-II was pre-treated for 2 h with NanH and/or BgaA and then incubated overnight with CMP-Neu5biotin (**1**) and immunoblotted with streptavidin-800CW. **b)** Cst-II was oxidized with 1 mM $NaIO_4$ and then incubated overnight with CMP-Neu5biotin or (**c**) CMP-Neu5biotin and fetuin.

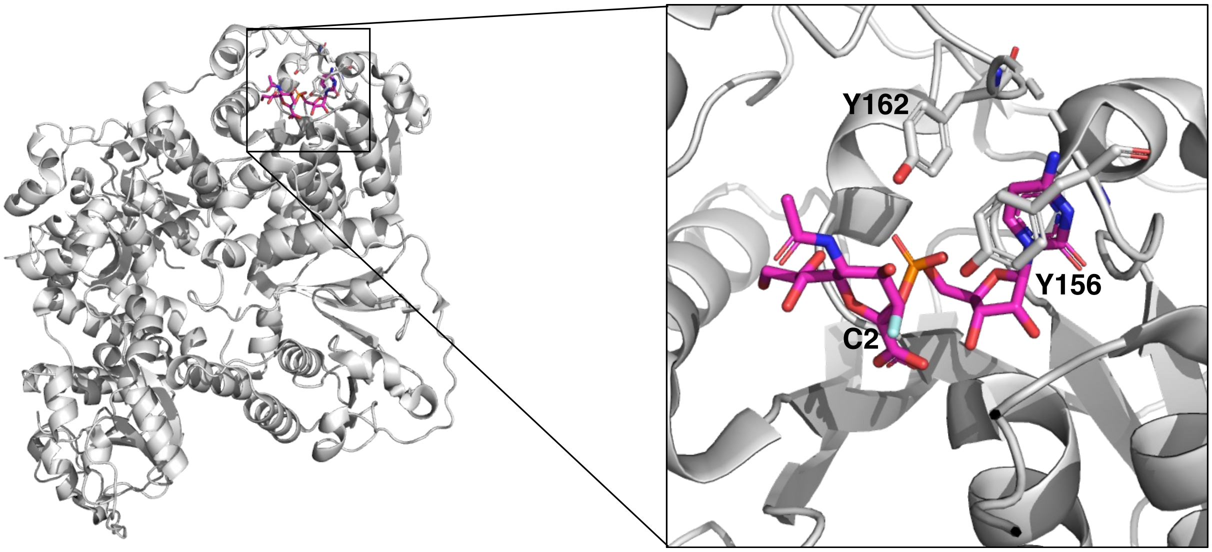


**Figure S9. Organization of the Cst-II active site.** The crystal structure of Cst-II in complex with donor analog CMP-3FNeu5Ac reported by Chiu et al. is represented (PDB = 1RO7) as visualized in PyMOL.[1] Tyr residues in the active site (Y156 and Y162) are shown. The substrate CMP-3FNeu5Ac is highlighted in purple with the anomeric carbon labeled (C2).

#### 2. Materials and Methods

Biotinylated *Sambucus nigra* agglutinin (SNA) (cat # B1305), biotinylated *Maackia amurensis* lectin-II (MAL-II) (cat # B1265), and SNA-FITC (cat # FL-1301-2) were purchased from Vector labs. Streptavidin-PacificBlue conjugate (cat # S11222) was purchased from Thermo-Fisher Scientific. Revert Revert™ 700 Total Protein Stain (cat # 926-11021) and Streptavidin-IRDye® 800CW (cat # 926-32230) were purchased from Li-Cor Biosciences.

**Sialidase immobilization and preparation of asialofetuin (AFet)**

NHS-activated agarose resin was purchased from Thermo-Fisher Scientific (cat # 26196). Enzyme immobilization was carried out following the manufacturer's instructions. A mixture of NHS-activated agarose resin (30 mg) and *C. perfringens* sialidase (NanH) in PBS pH 7.4 (0.75 mL, 15 mg) was incubated at room temperature for 1 h, inverting gently. The resin was then washed with 0.8 mL of PBS three times by centrifugation, and the supernatant was collected to measure the concentration of free enzyme that was not immobilized. Subsequently, 0.4 mL of quench buffer (1 M Tris-HCl, pH 7.4) was added to the resin to block remaining active NHS sites. The reaction was mixed for 20 min at room temperature, inverting gently. After completion, the resin was washed three times with 0.8 mL of PBS and kept in PBS with 20% glycerol at -80°C for a long-term storage. The amount of total protein in the collected wash supernatant was quantified by BCA assay (Pierce, Thermo-Fisher). Approximately 14.0 mg enzyme was immobilized (94%).

For the preparation of asialofetuin, bovine fetuin (Sigma, cat # F2379) was suspended in PBS (1 mg/mL) and incubated with immobilized NanH (0.1 μg/μg glycoprotein substrate) overnight at 37 °C, rocking gently. The reaction was then gently centrifuged, and the supernatant was recovered from the beads for subsequent use.

**Fetuin glycan labeling**

Fetuin or asialofetuin (10 μg) was suspended in PBS (10 μL) containing the CMP-Neu5Ac derivative (30 equivalents), with either ST6Gal1 (0.05 μg/μg glycoprotein), ST3Gal1 (0.2 μg/μg glycoprotein), or Cst-II (0.25 μg/μg glycoprotein) at a final concentration of 1 mg/mL fetuin/asialofetuin. Reactions were incubated at 37 °C, rocking gently overnight. Following incubation, samples were stored at −80 °C until analysis by immunoblotting.

**Cell culture**

RAJI and U937 cells were cultured in RPMI-1640 medium (with L-glutamine, sodium bicarbonate) supplemented with 10% fetal bovine serum (FBS) and 1X penicillin/streptomycin (P/S). Cells were passaged after reaching $2 \times 10^6$ viable cells/mL. SK-BR-3, HS-578-T, and HEK293T/17 cells were cultured in high glucose DMEM medium with L-glutamine, supplemented with 10% FBS and 1X P/S. MCF-7 cells were cultured in MEM with Earle's salts, L-glutamine, and sodium bicarbonate supplemented with 10% FBS and 1X P/S. Adherent cells were passaged using 0.25% trypsin-EDTA after reaching ~80% confluency. All cells were maintained in a humid 5% $CO_2$ atmosphere at 37 °C.

HEK293T MGAT1 KO cells were generated by CRISPR/Cas9. Briefly, MGAT1 targeting gRNA was cloned into lentiCRISPRv2 plasmid (Dr. Zhang, Addgene cat # 52961) using synthetic oligos (5'-CACCGGCCCGACCTGAGCAGCATTG-3'; 5'-AAACCAATGCTGCTCAGGTCGGGCC-3'). The cloned construct was packaged into lentiviral particles by transfection in HEK293T/17 cells (ATCC, cat # CRL-11268) along with psPAX2 (Dr. Didier Trono, Addgene, cat # 12260) and VSV-G (Dr. Bob Weinberg, Addgene, cat # 8454). Subsequently, HEK293T/17 cells were transduced and MGAT1 KO cells were selected for with 1 μg/mL puromycin. The bulk population was single-cell cloned by limiting dilution, and MGAT1 expression of clonal populations was determined by Western blotting using a rabbit monoclonal MGAT1 antibody (Abcam, cat # ab180578) with a goat anti-rabbit IgG Alexa Fluor® 488 Conjugate (NEB, cat # 4412S) and visualized on a LICOR Odyssey CLx. Anti-β-actin (Abcam, cat # ab8226) was used as a loading control. The selected clone was further screened by flow cytometry, staining with SNA-FITC.

**Glycosylation inhibitor treatments**

HS-578-T cells were plated in 12-well plates (100 000 cells/well) and incubated at 37 °C in 1 mL of DMEM medium containing 10% FBS and 1X P/S with either kifunensine (10 μg/mL, Carbosynth), benzyl-α-GalNAc (2.0 mM, Sigma), or GENZ-123346 (5 μM, Abcam) for 72 h.

**Sialidase treatment**

Cells were washed three times in serum-free culture medium, then treated as previously described.[2] Washed cells were incubated with 50 μg/mL *Clostridium perfringens* α2,3/6/8 sialidase (NanH) or 50 μg/mL *Streptococcus pneumoniae* α2,3-specific sialidase (Neu-S) in serum-free culture medium for 30 min and then were washed three times with serum-free culture medium prior to subsequent treatment.

**Cell-surface glycan labeling with Cst-II**

Adherent cells were plated in 12-well plates (300 000 cells/well) and grown to 80% confluency. Suspended cells were centrifuged and resuspended at 6 M cells/mL (100 μL per treatment). Prior to labeling, cells were washed three times with culture medium without FBS. Washed cells were incubated with a mixture of serum-free culture medium containing 100 μM CMP-Neu5Ac derivative, 0.1% BSA, and 200 μg/mL Cst-II for 2 h at 37 °C. Untreated control experiments were treated with a mixture of serum-free culture medium containing 0.1% BSA with or without Cst-II or CMP-Neu5Ac derivative. Following glycan labeling, cells were washed three times with 1% FBS/DPBS then treated as indicated.

For two-step labeling, cells were washed in DPBS and then incubated with 50 μM DBCO-biotin (Sigma, cat # 760706) in DPBS for 2 h at room temperature. Cells were washed three times with 1% FBS/DPBS then treated as indicated.

**Flow cytometry analysis**

For the detection of cell-surface α-2,6-linked or α-2,3-linked sialic acid, control and sialidase treated adherent cells were first washed three times with DPBS without Ca/Mg and detached using 10 mM EDTA for 5 min at 37 °C. Cells were then suspended in 1% FBS/DPBS, centrifuged gently (300 rpm for 3 min), washed twice with 1% FBS/DPBS, and resuspended in 200 µL of staining buffer (1% FBS/DPBS) with 20 µg/mL biotinylated-SNA or biotinylated-MAL-II for 30 min at 4 °C. Cells were then washed/centrifuged once with staining buffer, then incubated with streptavidin-Pacific Blue (2.5 µg/mL in staining buffer) for 30 min at 4 °C in the dark. The cells were washed once in staining buffer, centrifuged gently, and resuspended in 300 µL of FACS buffer (PBS without Ca/Mg supplemented with 2 mM EDTA and 0.5% BSA) and transferred to 96-well plates for flow cytometric analysis (Beckman Coulter, Cytoflex S). Cell viability was determined by adding 1 µg/mL PI to cell suspensions 1 min prior to analysis. The live population of cells was gated based on forward and side scatter emission, and exclusion of PI positive cells on the FL11 (610/20 BP filter) emission channel. Streptavidin-Pacific Blue binding was determined by fluorescence intensity on the FL6(450/45 BP filter) emission channel.

For the detection of cell-surface glycan biotinylation using streptavidin, enzymatically labeled cells were stained in-well with streptavidin-Pacific Blue (2.5 µg/mL, Thermofisher) in 1% FBS/DPBS for 30 min at 4 °C in the dark. The cells were washed three times with DPBS without Ca/Mg, and detached using 10 mM EDTA for 5 min at 37 °C. The cells were washed in 1% FBS/DPBS, centrifuged gently (300 rcf, 3 min), and resuspended in 300 µL of FACS buffer (PBS without Ca/Mg supplemented with 2 mM EDTA and 0.5% BSA) and transferred to 96-well plates for flow cytometric analysis (Beckman Coulter, Cytoflex S). Cell viability was determined by adding 1 µg/mL PI to cell suspensions 1 min prior to analysis. The live population of cells was gated based on forward and side scatter emission, and exclusion of PI positive cells on the FL11 (610/20 BP filter) emission channel. Streptavidin-Pacific Blue binding was determined by fluorescence intensity on the FL6 (450/45 BP filter) emission channel. Data points are representative of at least 3 separate experiments (n = 3).

**PNGase F treatment**

Fetuin or cell lysates enzymatically labeled as described above were treated with PNGase F to remove N-glycans according to the manufacturer's protocol (NEB cat # P0704). Proteins were denatured with 1X glycoprotein denaturing buffer by incubation at 100 °C for 10 min. After chilling on ice for 10 min, samples were adjusted to contain 1X GlycoBuffer 2 and 1% NP-40 prior to addition of PNGase F (2 U/µg total protein) and overnight incubation rocking at 37 °C. Untreated control experiments were treated identically excluding the addition of PNGase F.

**Immunoblot analysis**

Following enzymatic cell-surface glycan labeling, cells were washed with cold DPBS and lysed with RIPA lysis buffer with 1X protease inhibitor cocktail (NEB). The cell lysates were clarified by centrifugation at 14 000g for 15 min and the total protein content of the clear supernatants was quantified by BCA assay. The samples (30 µg total protein) were resolved on an 8% SDS-PAGE gel and transferred to a low-fluorescence PVDF membrane (Immobilon-FL, Sigma). For immunoblotting of fetuin, samples (2 µg protein) were resolved on 10% SDS-PAGE gels. For normalization, total protein was stained with Revert Total Protein stain (Li-Cor Biosciences) for 5 min, washed twice with wash buffer (6.7% acetic acid, 30% methanol), and rinsed briefly in TBS prior to scanning on an Odyssey Li-Cor CLx scanner (Li-Cor Biosciences).

Next, the membrane was blocked in blocking buffer (5% nonfat dry milk in TBS) for 1 h at room temperature. The blocked membrane was subsequently incubated for 1 h at room temperature with a fluorescent streptavidin-800CW conjugate (Li-Cor Biosciences) in blocking buffer with 0.1% Tween-20 and washed with TBST (4 × 5 min). The membrane was then rinsed in TBS prior to detection of fluorescence by scanning on an Odyssey Li-Cor CLx scanner (Li-Cor Biosciences).

**Detection of cell-surface labeling by fluorescence microscopy**

HS578T cells were plated in 12-well plates (150 000 cells/well) and were grown to ~60% confluency then labeled as described above with a total volume of 200 µL. Cells labeled in one step with CMP-Neu5biotin (**1**) were washed three times in 1% FBS/DPBS and stained with avidin-AlexaFluor-488 (2.5 µg/mL) in 1% FBS/DPBS for 30 min at 4 °C in the dark. Cells labeled in two steps with CMP-Neu5Az (**2**) were washed three times in DPBS and incubated with 10 µM DBCO-AZDye488 (Click Chemistry Tools, cat # 1278-1) in DPBS for 1 hour at room temperature in the dark. After washing three times in 1% FBS/DPBS, cells incubated with DBCO-s-biotin were stained with avidin-AlexaFluor-488 (2.5 µg/mL) in 1% FBS/DPBS for 30 min at 4 °C in the dark. Cells were washed with 1% FBS/DPBS and fixed with formaldehyde (3.7% in PBS) at room temperature for 15 min, then washed and permeabilized for 10 min at room temperature using 0.1% Triton X-100 in DPBS. After washing, the nuclei were labeled with 0.3 µM DAPI in PBS for 30 min before washing in DPBS and imaging by fluorescent microscopy using a 20× air objective.

**Assessment of CMP-Neu5biotin hydrolysis with Cst-II**

0.5 mg CMP-Neu5biotin (**1**) was dissolved in Tris-HCl buffer (100 mM, pH 9) to a final concentration of 10 mM, with 20 mM $MgCl_2$. Then Cst-II (100 µg/ml) was added to the reaction tube and the reaction was shaken at 180 rpm and 37°C. After shaking for 2 hours the enzyme was removed by centrifugation using an Amicon Ultra-10 (MWCO-10k) centrifugal filter. The filtrate was lyophilized and analyzed by NMR (700 MHz, $D_2O$ referenced to 4.79 ppm).

**Assessment of Cst-II self-labeling**

For the glycosidase pre-treatment of Cst-II, Cst-II (10 µg) was suspended in PBS and incubated with or without the addition of NanH (0.2 µg/µg substrate) and/or BgaA (0.5 µg/µg substrate) rocking gently for 2 h at 37 °C. Biotinylated CMP-Neu5Ac derivative **1** (30 equivalents) was then added, and the reaction was rocked gently overnight at 37 °C. Following incubation, samples were stored at −80 °C until analysis by immunoblotting.

For the periodate oxidation on Cst-II, protocols were modified from those previously described.[3] Cst-II (100 µg) was buffer exchanged into PBS using a 10 kDa MWCO spin filter (Amicon). A 10 µg aliquot of Cst-II was then treated with sodium periodate at a final concentration of 1 mM, which was incubated on ice, in the dark for 30 min. The reaction was quenched by the addition of 1 mM glycerol, buffer exchanged into PBS, and total protein was quantified by BCA assay. CMP-Neu5Ac derivative (30 equivalents) was then added, and the reaction was rocked gently overnight at 37 °C. Following incubation, samples were stored at −80 °C until analysis by immunoblotting.

#### *Bacterial protein expression*

Bacterial proteins *P. multocida* strain Pm70 inorganic pyrophosphatase (PmPpA), *E. coli* K12 sialic acid aldolase (aldolase), *N. meningitidis* CMP-sialic acid synthetase (CSS), *C. perfringens* α2,3/6/8 sialidase (NanH), *S. pneumoniae* α2,3 sialidase (Neu-S), *S. pneumoniae* β-galactosidase (BgaA), metagenomic alkaline phosphatase (mAP), and *C. jejuni* sialyltransferase II (Cst-II) were recombinantly expressed as previously reported.[2, 4-9] The genes encoding aldolase and PmPpA were commercially synthesized and inserted into pET-15b vectors using NcoI and BamHI or NdeI and BamHI restriction sites, respectively (Genscript). The gene encoding *N. meningitidis* CMP-sialic acid synthetase (CSS) was commercially synthesized and inserted into a pET-22b(+) vector using NdeI and XhoI restriction sites (Genscript). The gene sequence of *C. perfringens* a2,3/6/8 neuraminidase (NanH) (GenBank: Y00963.1) was commercially synthesized and ligated into a pET-15b plasmid using NdeI and XhoI restriction sites (Genscript). The gene encoding metagenomic alkaline phosphatase (mAP) was commercially synthesized and ligated into a pET-21a(+) plasmid using NdeI and XhoI restriction sites (Genscript). The gene encoding a truncated mutant of Cst-II having a C-terminal 32-amino-acid deletion and a single mutation I53S was commercially synthesized into a pET-22b(+) vector (Genscript). The plasmids encoding the genes for Neu-S and BgaA were generously provided by Dr. Matthew Macauley (University of Alberta) and Dr. David Kwan (Concordia University), respectively.

**General Expression of Bacterial Proteins**

The vectors containing the genes of interest were chemically transformed into BL-21 competent cells. Transformed cells were cultured in 5 mL LB media supplemented with 100 μg/mL ampicillin (LB-amp) for 16 h at 37 °C with shaking (200 rpm) for 16 hours. Cultured cells were then scaled up to 500 mL in LB-amp to an optical density ($OD_{600}$) of 0.8. Protein expression was then induced with addition of isopropyl β-D-thiogalactopyranoside (IPTG) to a final concentration of 0.1 mM and cells were incubated at 25 °C with shaking (200 rpm) for 16 hours before harvesting by centrifugation. Pelleted cells were resuspended in lysis buffer (0.1 M Tris-HCl, pH 8.0, 0.1% Triton X-100) and lysed by high pressure homogenization using an Avestin Emulsiflex C3. Cell debris was removed by ultracentrifugation, and the supernatant was mixed 1:1 with binding buffer (10mM imidazole, 500mM NaCl, 50mM Tris-HCl, pH 7.5).

**Expression of BgaA**

The vector containing the gene for BgaA was chemically transformed into BL-21 competent cells. Transformed cells were cultured in 5 mL LB media supplemented with 50 μg/mL kanamycin (LB-kanamycin) for 16 h at 37 °C with shaking (200 rpm) for 16 hours. Cultured cells were then scaled up to 500 mL in LB-kanamycin to an optical density ($OD_{600}$) of 0.6. Protein expression was then induced with addition of isopropyl β-D-thiogalactopyranoside (IPTG) to a final concentration of 0.5 mM and cells were incubated at 20 °C with shaking (200 rpm) for 16 hours before harvesting by centrifugation. Pelleted cells were resuspended in lysis buffer (0.1 M Tris-HCl, pH 8.0, 0.1% Triton X-100) and lysed by high pressure homogenization using an Avestin Emulsiflex C3. Cell debris was removed by ultracentrifugation, and the supernatant was mixed 1:1 with binding buffer (10mM imidazole, 500mM NaCl, 50mM Tris-HCl, pH 7.5).

**Expression of mAP**

The vector containing the gene for mAP was chemically transformed into BL-21 competent cells. Transformed cells were cultured in 5 mL LB media supplemented with 100 μg/mL ampicillin for 16 h at 37 °C with shaking (200 rpm) for 16 hours. Cultured cells were then scaled up to 500 mL in LB-amp to an optical density ($OD_{600}$) of 0.4. Protein expression was then induced with addition of isopropyl β-D-thiogalactopyranoside (IPTG) to a final concentration of 0.5 mM and cells were incubated at 15 °C with shaking (200 rpm) for 40 hours before harvesting by centrifugation. Pelleted cells were resuspended in lysis buffer (0.1 M Tris-HCl, pH 8.0, 0.1% Triton X-100) and lysed by high pressure homogenization using an Avestin Emulsiflex C3. Cell debris was removed by ultracentrifugation, and the supernatant was mixed 1:1 with binding buffer (10mM imidazole, 500mM NaCl, 50mM Tris-HCl, pH 7.5).

**Bacterial Protein Purification**

The resulting solutions were then filtered using a 0.22 μm Filter Unit and applied to a $Ni^{2+}$-sepharose resin (GE Healthcare, fast flow) pre-equilibrated with binding buffer. The column was washed with 6 column volumes (CV) of binding buffer, 6 CV wash buffer (50mM imidazole, 500mM NaCl, 50mM Tris–HCl, pH 7.5), and eluted with 10 CV elution buffer (200mM imidazole, 500mM NaCl, 50mM Tris-HCl, pH 7.5). Overexpression and purification were assessed by SDS-PAGE. Purified protein was buffer exchanged into 20 mM Tris-HCl (pH 7.5) buffer and concentrated using a 10 kDa MWCO spin filter (Amicon). All proteins were stored in 20 mM Tris-HCl (pH 7.5) with 20% glycerol at -80 °C, except for CSS which was stored in 20 mM Tris-HCl (pH 8.5) with 20% glycerol at -80 °C, and BgaA which was stored in 20 mM Tris (pH 7.5) containing 100 mM NaCl. Final protein concentrations were determined by BCA assay. Yields per 500 mL culture were: aldolase = 40 mg; CSS = 45 mg; PmPpA = 55 mg; NanH = 175 mg; Neu-S = 22 mg; BgaA = 151 mg; mAP = 5 mg; Cst-II = 102 mg.

#### *Mammalian protein expression*

The plasmids encoding soluble, secreted GFP fusion proteins containing the catalytic domain of human β-galactoside α-2,6-sialyltransferase 1 (ST6Gal1) or β-galactoside α-2,3-sialyltransferase 1 (ST3Gal1) in pGEn2-DEST vectors were purchased from DNASU (cat # HsCD00413052, HsCD00413169). Recombinant GFP-ST6Gal1 and GFP-ST3Gal1 protein was expressed and purified as previously described using Expi293 cells.[2, 10] Plasmids were chemically transformed into DH5-alpha competent cells. Transformed cells were cultured in 5 mL LB media supplemented with 100 μg/mL ampicillin for 16 h at 37 °C and shaking (200 rpm) for 16 hours. Cultured cells were then scaled up to 500 mL and plasmids were isolated using the PureLink™ HiPure Plasmid Maxiprep Kit (Thermofisher, cat # K210006).

**Expi293 cell maintenance**

Expi293 cells (ThermoFisher) were cultured in Expi293 Expression Medium (ThermoFisher). Cells were maintained in a humid 5% $CO_2$ atmosphere at 37 °C with shaking at 120 rpm. Cells were passaged for maintenance after reaching $4 \times 10^6$ viable cells/mL. Cells were cultured for at least 3 passages following thaw prior to transfection.

**Transfection and Expression**

Expi293 cells were transiently transfected with the vector containing the ST6Gal1 or ST3Gal1 gene using the Expifectamine 293 Transfection Kit (ThermoFisher, cat # A14524). On day 5 following transfection, cells were harvested by centrifugation (25 min, 4000 rcf). The supernatant was collected and adjusted to contain 20 mM imidazole, 200 mM NaCl, and 30 mM sodium phosphate, at pH 7.2. The resulting solutions were then filtered using a 0.45 µm PES Filter Unit and applied to a $Ni^{2+}$-sepharose resin pre-equilibrated with column buffer (20 mM HEPES, 300 mM NaCl, pH 7.2) containing 20 mM imidazole. The column was sequentially washed with column buffer containing 20 mM, 50 mM, and 100 mM imidazole prior to elution with 300 mM imidazole, and the column was flushed with 500 mM. Overexpression and purification were assessed by SDS-PAGE. Purified protein was buffer exchanged into 20 mM Tris-HCl (pH 7.5) buffer, concentrated using a 10 kDa MWCO spin filter (Amicon), and stored in 20 mM Tris-HCl (pH 7.5) with 20% glycerol at -80 °C. The final protein concentration was determined by BCA assay. Yields per 100 mL culture were: ST6Gal1 = 4.5 mg; ST3Gal1 = 4.1 mg.

#### Chemical and Enzymatic Synthesis

##### Propargyl biotinamide (S1)

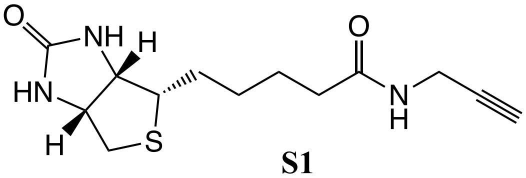


**S1**

D-Biotin (361 mg, 1.48 mmol) was dissolved in 14.6 mL dry MeCN/MeOH solution (3:1 v/v) under Ar. Propargylamine (114 µL, 1.78 mmol) and EDC•HCl (447 mg, 2.33 mmol) were added, and the mixture was stirred overnight. The reaction mixture was concentrated *in vacuo* and the residue was resuspended in cold MeOH and filtered through Celite. The filtrate was concentrated *in vacuo* and purification by flash column chromatography (9:1 $CH_2Cl_2$/MeOH) afforded the product as a white powder (120 mg, 29%). $^{1}H$ NMR (500 MHz, $D_2O$) δ 4.53 (dd, J = 7.8, 4.8 Hz, 1H, $CH_2C\mathit{H}NH$), 4.35 (dd, J = 8.0, 4.5 Hz, 1H, $CHC\mathit{H}NH$), 3.87 (s, 2H, $NHC\mathit{H_2}C$), 3.26 (m, 1H, CHS), 2.92 (dd, J = 13.0, 5.0 Hz, 1H $CH_2S$), 2.70 (d, J = 13.0 Hz, 1H, $CH_2$'S), 2.20 (t, J = 7.3 Hz, 2H, $CH_2CH_2CH_2C\mathit{H_2}C{=}O$), 1.69 – 1.47 (m, 4H, $C\mathit{H_2}CH_2C\mathit{H_2}CH_2C{=}O$), 1.39 – 1.29 (m, 2H, $CH_2C\mathit{H_2}CH_2CH_2C{=}O$). $^{13}C$ NMR (176 MHz, $D_2O$) δ 176.80, 62.02, 60.24, 55.28, 39.66, 35.24, 28.63, 27.72, 27.57, 24.95. HRMS (ESI-MS) m/z calculated for *m/z* $[M+H]^+$ cald. for $C_{13}H_{20}N_3O_2S^+$: 282.1276, found: 282.1271.

##### CMP-Neu5Az (2)

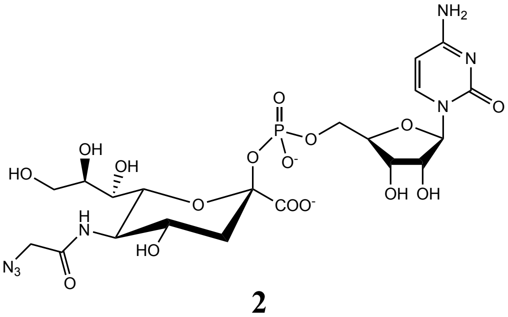


**2**

ManNAz was prepared as previously described in the literature.[11, 12] ManNAz (20 mg, 0.095 mmol), sodium pyruvate (52.46 mg, 0.48 mmol), and CTP disodium salt (65.33 mg, 0.12 mmol) were dissolved in a Tris-HCl buffer (0.1 M, pH 8.8) containing $MgCl_2$ (20 mM) to a final ManNAz substrate concentration of 50 mM. To this mixture, recombinant *E. coli* aldolase (35 mg/mmol substrate), *N. meningitidis* CMP-sialic acid synthetase (31 mg/mmol substrate) and *P. multocida* inorganic pyrophosphatase (18 mg/mmol substrate) were added and the reaction was incubated at 37 °C with shaking (200 rpm), in the dark. Reaction progress was monitored by TLC (7:3 EtOH/1 M aqueous ammonium bicarbonate), which was allowed to proceed until disappearance of the ManNAz starting material (Rf = 0.8), detected by staining with *p*-anisaldehyde. The reaction was then quenched by the addition of metagenomic alkaline phosphatase (mAP) to degrade unreacted CTP to aid subsequent purification steps. The mixture with mAP was incubated at 37 °C with shaking until CTP was no longer visible by TLC (detected by UV exposure; Rf = 0.1), which otherwise co-elutes with the product over P2-BioGel purification. Precipitates were removed by centrifugation and enzymes were then removed by spin-filtration (10 kDa MWCO, Amicon Ultra-15 Centrifugal Filter, cat # UFC9010) before flash-freezing with liquid $N_2$ and lyophilization. The isolated residue was resolubilized in 0.1 M ammonium bicarbonate buffer and loaded on a fine P2-BioGel® (BioRad) size exclusion column eluting with ice-cold 0.1 M $NH_4HCO_3$. Fractions containing the title compound were collected on ice, pooled, and lyophilized to afford CMP-Neu5Az (**2**) as a white powder (64 mg, 54%). CMP-Neu5Az (**2**) was stored as a dry solid at -80 °C and dissolved in an appropriate buffer immediately before use. Characterization data are consistent with previous reports.[4, 11] $^{1}$H NMR (700 MHz, $D_2O$) δ 7.98 (d, *J* = 7.6 Hz, 1H, $H6^{Cyt}$), 6.13 (d, *J* = 7.6 Hz, 1H, $H5^{Cyt}$), 6.00 (d, *J* = 4.6 Hz, 1H, $H1^{Rib}$), 4.35 (t, *J* = 4.9 Hz, 1H, $H3^{Rib}$), 4.31 (t, *J* = 4.9 Hz, 1H, $H2^{Rib}$), 4.25-4.23 (m, 4H, $H4^{Rib}$, $H5^{Rib}$, $H5'^{Rib}$, $H6^{Sia}$), 4.16-4.11 (m, 1H, $H4^{Sia}$), 4.09 (s, 2H, CO-$CH_2$-$N_3$), 4.03 (t, *J* = 10.3 Hz, 1H, $H5^{Sia}$), 3.94 (ddd, *J* = 9.7 Hz, 6.5 Hz, 2.5 Hz, 1H, $H8^{Sia}$), 3.88 (dd, *J* = 11.8 Hz, 2.6 Hz, 1H, $H9^{Sia}$), 3.62 (dd, *J* = 11.8 Hz, 6.5 Hz, 1H, $H9'^{Sia}$), 3.45 (d, *J* = 9.7 Hz, 1H, $H7^{Sia}$), 2.51 (dd, *J* = 13.3 Hz, 4.8 Hz, 1H, $H_{3eq}{}^{Sia}$), 1.67 (ddd, *J* = 13.1 Hz, 11.9 Hz, 5.7 Hz, 1H, $H_{3ax}{}^{Sia}$). $^{13}$C-DEPT135 NMR (176 MHz, $D_2O$) δ 141.65 ($C6^{Cyt}$), 96.81 ($C5^{Cyt}$), 88.96 ($C1^{Rib}$), 82.91 ($C4^{Rib}$), 74.29 ($C2^{Rib}$), 71.31 ($C6^{Sia}$), 69.67 ($C8^{Sia}$), 69.26 ($C3^{Rib}$), 68.80 ($C7^{Sia}$), 66.60 ($C4^{Sia}$), 64.80 ($C5^{Rib}$), 62.87 ($C9^{Sia}$), 51.98 ($C5^{Sia}$), 51.92 (CO-$CH_2$-$N_3$), 41.16 ($C3^{Sia}$). ESI-MS m/z calcd for $C_{20}H_{29}N_7O_{16}^-$, $[M-H]^-$ : 654.141, found: 654.144; m/z calculated for *m/z* $[M+Na]^-$ calcd for $C_{20}H_{28}N_7O_{16}PNa^-$: 676.123, found: 676.126.

**CMP-Neu5biotin (1)**

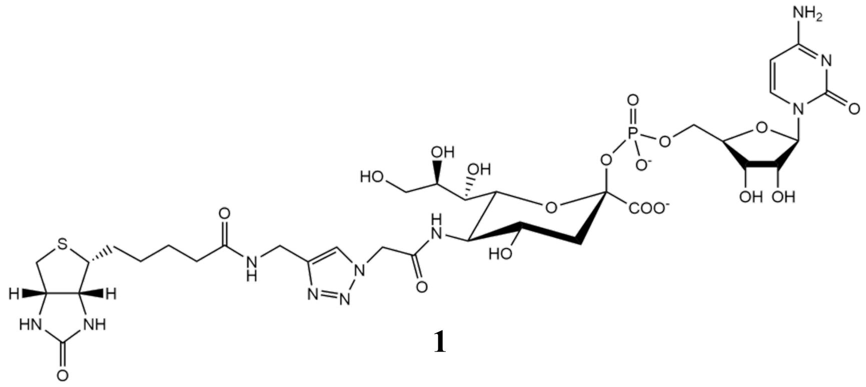


**1**

CMP-Neu5biotin (**1**) was prepared based on protocols described in the literature.[11, 12] Stock solutions of 0.1 M CuSO-4, 0.2 M sodium L-ascorbate, and 0.1 M THPTA in 0.1 M $NH_4HCO_3$ were freshly made before each Cu-catalyzed azide-alkyne cycloaddition (CuAAC) reaction. To a solution of CMP-Neu5Az (5 mg, 7.6 µmol) and propargyl biotinamide (2.7 mg, 9.5 µmol) in 300 uL 0.1 M $NH_4HCO_3$, 0.1 M CuSO4 (76.3 µL, 7.6 µmol), 0.1 M THPTA (38.1 µL, 3.8 µmol), and 0.2 M sodium L-ascorbate (38.1 µL, 11.4 µmol) were added. The resulting mixture was stirred at room temperature for 3 hours to have minimal hydrolysis of the product. The mixture was then directly loaded onto a fine P2-BioGel® (BioRad) size exclusion column eluting with ice-cold 0.1 M $NH_4HCO_3$ and fractions were collected on ice. Fractions containing the product were identified by spot test to analyze by NMR, combined, and lyophilized to afford the title compound as a white powder (6.8 mg, 68.3%). CMP-Neu5biotin was stored as a dry solid at -80 °C and was dissolved in an appropriate buffer immediately before use. $^{1}$H NMR (700 MHz, $D_2O$) δ 8.08 (d, J = 7.7 Hz, 1H, H-6 cyt), 7.98 (s, 1H, CH=C, triazole), 6.21 (d, J = 8.4 Hz, 1H, H-5 cyt), 5.99 (d, J = 4.2 Hz,

1H, H-1 rib), 5.33 (d, J = 16.8 Hz, 1H, triazole-$CH_2$-C=O), 5.30 (d, J = 16.8 Hz, 1H, triazole-$CH_2$'-C=O), 4.62 (dd, J = 8.4, 4.9 Hz, 1H, $CH_2C\textit{H}NH$, biotin), 4.51 (d, J = 15.4 Hz, 1H, triazole-$CH_2$-NH) , 4.48 (d, J = 15.4 Hz, 1H, triazole-$CH_2$'-NH), 4.40 (dd, J = 8.1, 4.9 Hz, 1H, $CHC\textit{H}NH$, biotin), 4.36 (t, J = 4.9 Hz, 1H, H-3, rib), 4.33 (t, J = 4.9 Hz, 1H, H-2, rib), 4.28 – 4.24 (m, 3H, H-4 rib, H-5 rib), 4.23 (d, J = 9.8, 1H, H-6), 4.15 (td, J = 10.9, 4.9 Hz, 1H, H-4), 4.02 (t, J = 10.5 Hz, 1H, H-5), 3.94 (ddd, J = 9.5, 6.7, 2.1 Hz, 1H, H-8), 3.89 (dd, J = 12.3, 2.8 Hz, 1H, H-9a), 3.64 (dd, J = 11.6, 7.0, 1H, H-9b), 3.48 (d, J = 9.8, 1H, H7), 3.35 – 3.30 (m, 1H, CHS, biotin), 3.01 (dd, J = 13.3, 4.9 Hz, 1H $CH_2S$, biotin), 2.80 (d, J = 13.3 Hz, 1H, $CH_2$'S, biotin), 2.39 (dd, J = 13.3, 4.9 Hz, 1H, H-3eq), 2.31 (t, J = 7.0 Hz, 2H, $CH_2CH_2CH_2C\textit{H}_2C$=O, biotin), 1.72 – 1.55 (m, 5H, H-3ax + $C\textit{H}_2CH_2C\textit{H}_2CH_2C$=O, biotin), 1.40 – 1.35 (m, 2H, $CH_2C\textit{H}_2CH_2CH_2C$=O, biotin). $^{13}C$ NMR (176 MHz, $D_2O$) δ 142.4, 125.3, 96.2, 89.3, 83.0, 74.3, 69.7, 69.3, 64.8, 63.7, 62.9, 62.0, 60.2, 55.4, 52.1, 39.8, 35.2, 34.3, 27.6, 25.0. ESI-MS m/z calcd for $C_{33}H_{48}N_{10}O_{18}PS^-$, $[M-H]^-$ : 935.2612, found, 935.2652.

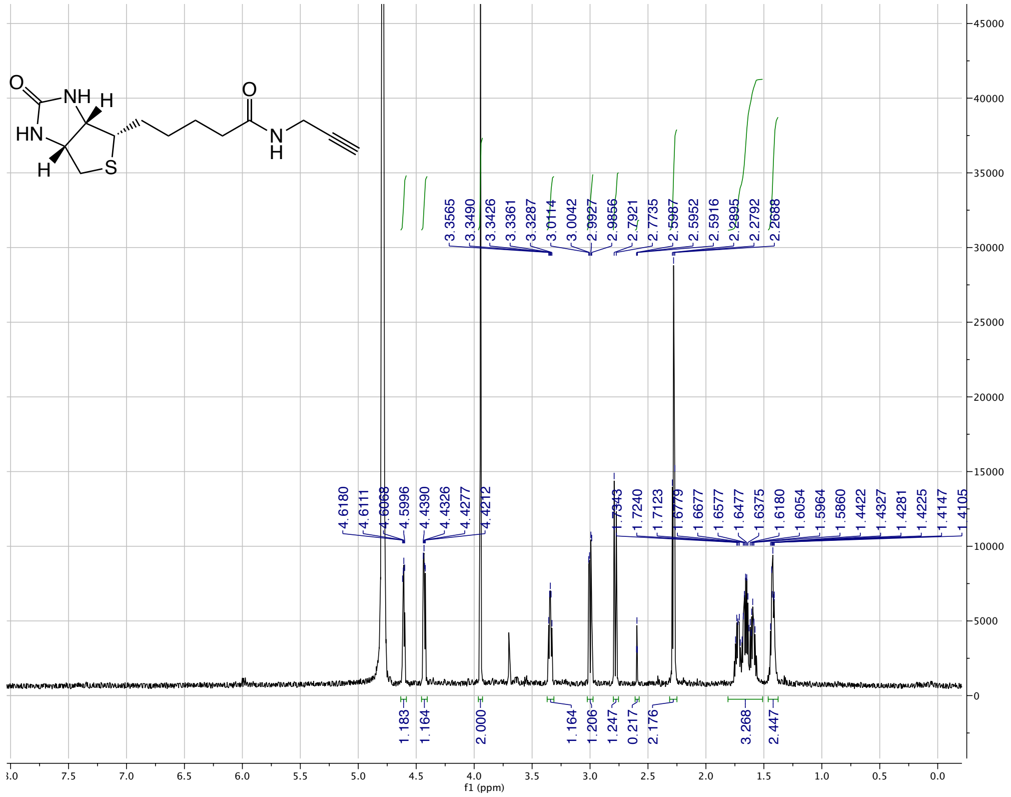
O
NH
H
HN
H
S
O
N
H
4.6180
4.6111
4.6068
4.5996
4.4390
4.4326
4.4277
4.4212
3.3565
3.3490
3.3426
3.3361
3.3287
3.0114
3.0042
2.9927
2.9856
2.7921
2.7735
2.5987
2.5952
2.5916
2.2895
2.2792
2.2688
1.7343
1.7240
1.7123
1.6779
1.6677
1.6577
1.6477
1.6375
1.6180
1.6054
1.5964
1.5860
1.4422
1.4327
1.4281
1.4225
1.4147
1.4105
1.183
1.164
2.000
1.164
1.206
1.247
0.217
2.176
3.268
2.447
f1 (ppm)

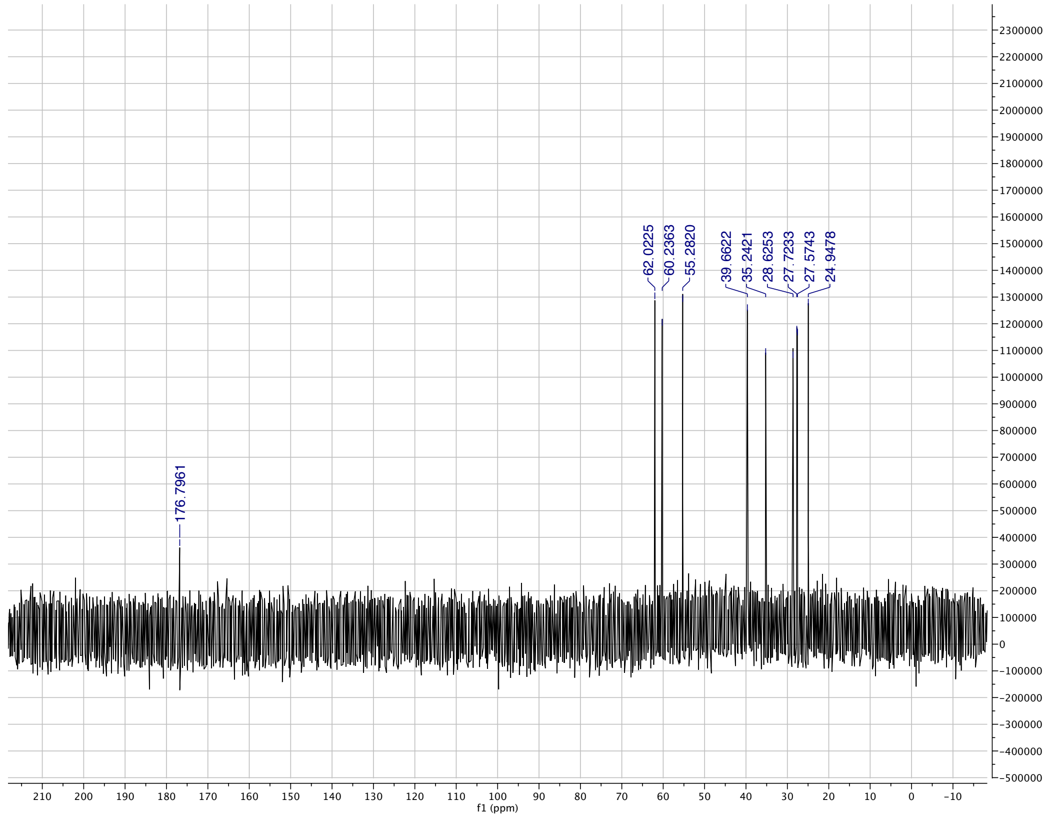
176.7961
62.0225
60.2363
55.2820
39.6622
35.2421
28.6253
27.7233
27.5743
24.9478
f1 (ppm)

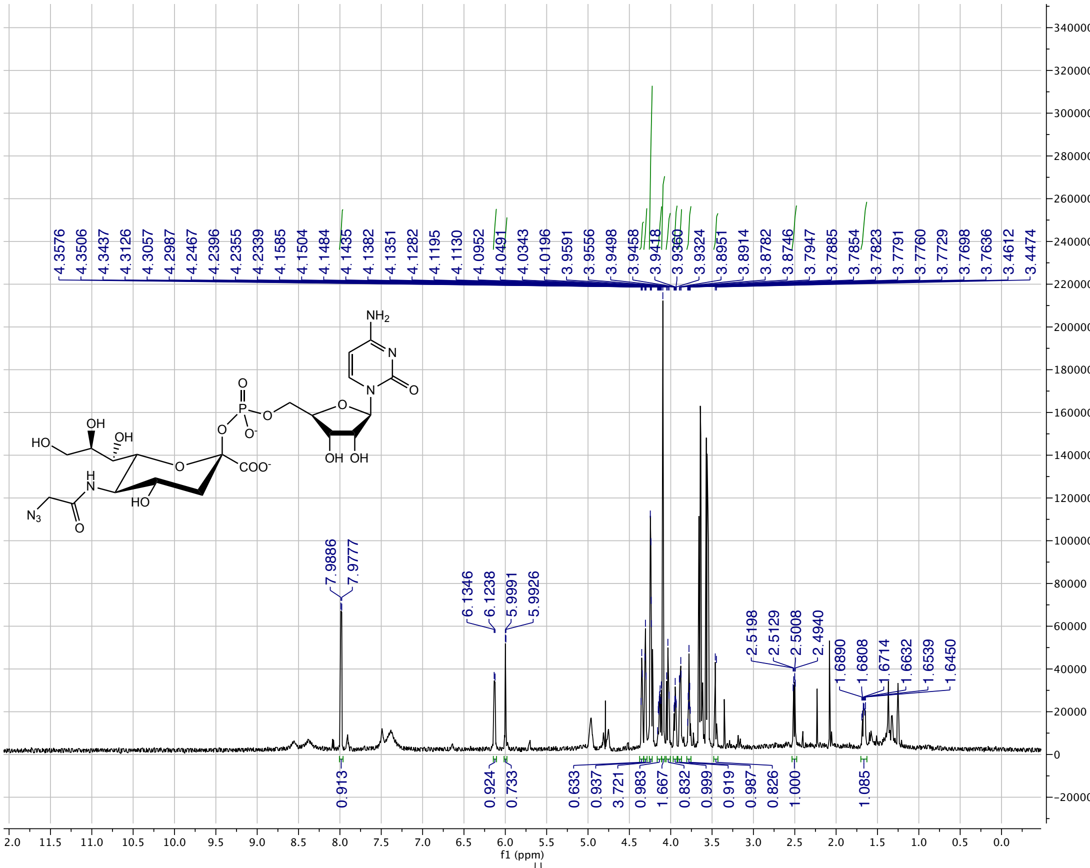
NH2
N
O
O
P
O
O
O
N
O
O-
OH
HO
OH
O
COO-
H
N
HO
OH
OH
N3
O
7.9886
7.9777
6.1346
6.1238
5.9991
5.9926
2.5198
2.5129
2.5008
2.4940
1.6890
1.6808
1.6714
1.6632
1.6539
1.6450
0.913
0.924
0.733
0.633
0.937
3.721
0.983
1.667
0.832
0.999
0.919
0.987
0.826
1.000
1.085
f1 (ppm)

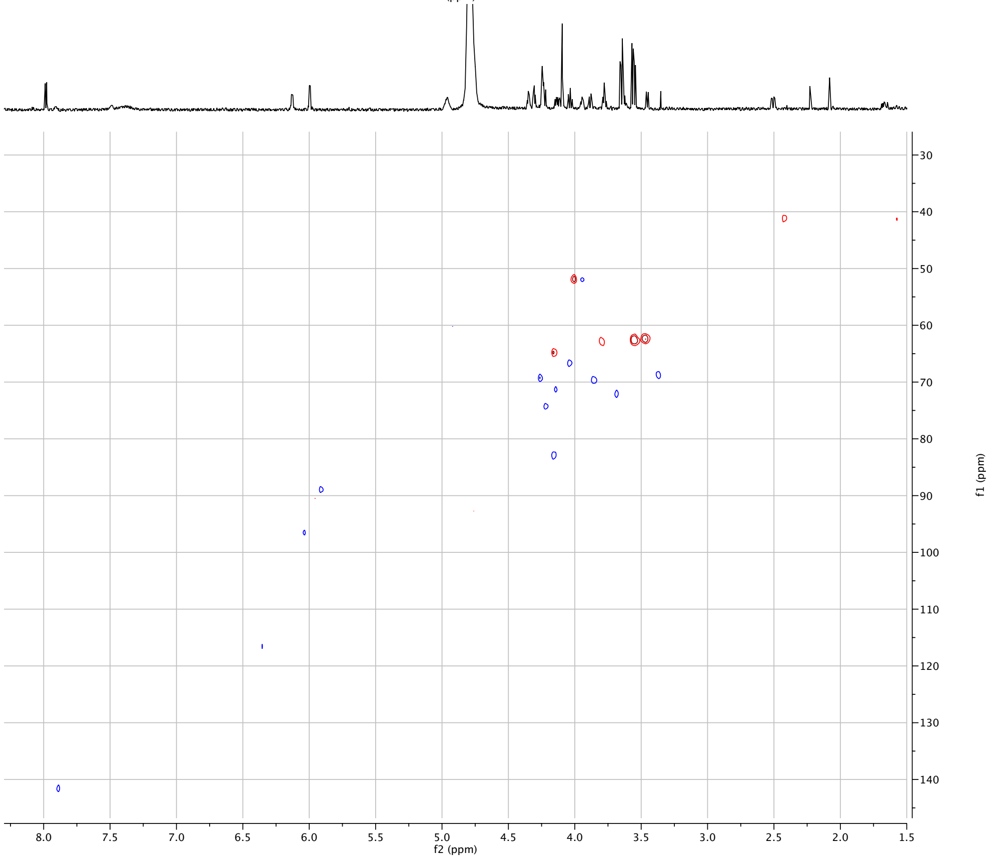
f2 (ppm)
f1 (ppm)

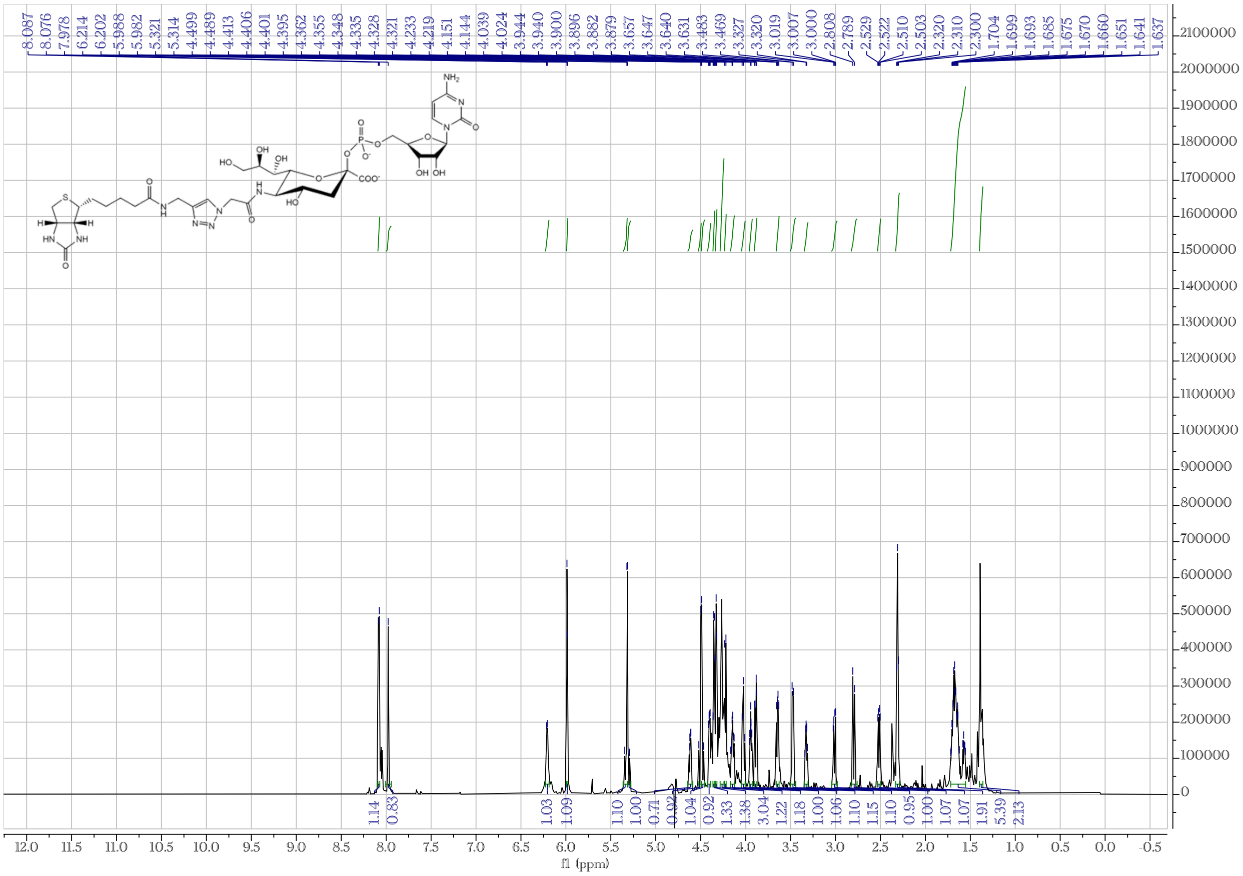
8.087 8.076 7.978 6.214 6.202 5.988 5.982 5.321 5.314 4.499 4.489 4.413 4.406 4.401 4.395 4.362 4.355 4.348 4.335 4.328 4.321 4.233 4.219 4.151 4.144 4.039 4.024 3.944 3.940 3.900 3.896 3.882 3.879 3.657 3.647 3.640 3.631 3.483 3.469 3.327 3.320 3.019 3.007 3.000 2.808 2.789 2.529 2.522 2.510 2.503 2.320 2.310 2.300 1.704 1.699 1.693 1.685 1.675 1.670 1.660 1.651 1.641 1.637
1.14 0.83 1.03 1.09 1.10 1.00 0.71 0.92 1.04 0.92 1.33 1.38 3.04 1.22 1.18 1.00 1.06 1.10 1.15 1.10 0.95 1.00 1.07 1.07 1.91 5.39 2.13
f1 (ppm)

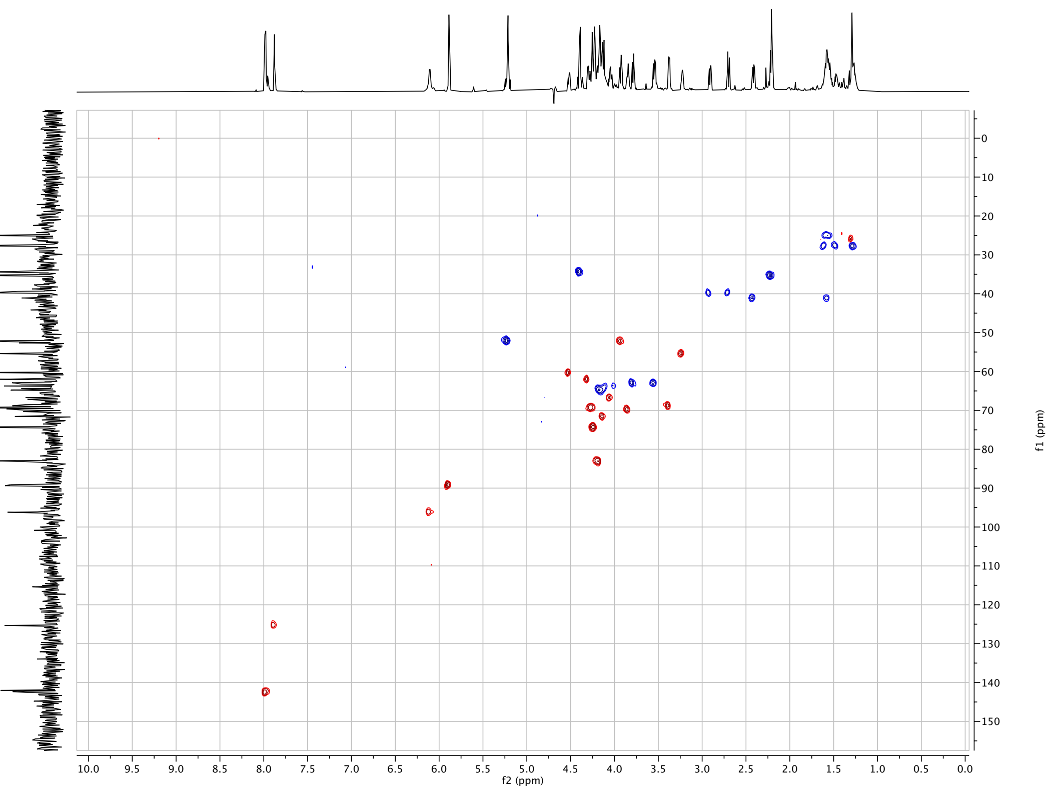
f2 (ppm)
f1 (ppm)
